# Genome-context-aware discovery of antibacterial peptides from bacterial small open reading frames

**DOI:** 10.64898/2026.08.17.745349

**Authors:** Qingxiu Li, Zhenjun Li

**Affiliations:** Institute of Infectious Diseases, Chinese Center for Disease Control and Prevention, 155 Changbai Road, Changping District, Beijing 102206, China

**Keywords:** small open reading frame, antimicrobial peptide, bacterial genomes, genome context, machine learning, peptide discovery

## Abstract

Small open reading frames (sORFs) are a potentially rich, yet error-prone, source of antimicrobial-peptide (AMP) candidates: short sequences are readily prioritized by AMP classifiers but may derive from incomplete gene calls. We developed a genome-context-aware discovery workflow that separates AMP-like sequence properties from evidence for a complete, recurrent coding locus. From 649,653 RefSeq assemblies representing 327 clinically relevant bacterial species, species-aware clustering and length filtering yielded 4,442,548 representative 10–100-aa sequences. AmpScanner v2, Macrel and AMPlify identified 585 non-haemolytic records supported by all three models. However, genome-context auditing of 11,918 mapped candidates showed that 529 of 536 mapped consensus candidates were supported exclusively by partial ORFs near contig termini. By contrast, 3,382 candidates had at least one complete non-edge occurrence; 1,069 recurred in ≥2 assemblies and 251 in ≥10 assemblies. We therefore assembled a 20-peptide panel through two explicitly labelled routes: sequence/structure-led selection (n=8) and genome-supported selection (n=12). Broth microdilution against Escherichia coli ATCC 25922 and Staphylococcus aureus ATCC 25923 identified low-micromolar activity in both routes. CAND_04141, a recurrent complete non-edge candidate, had the strongest combined profile (MICs of 4 and 2 μM, respectively), while CAND_07825 and CAND_04265 were also active at low micromolar concentrations. In plate-count MBC assays, all three advanced peptides achieved ≥3-log10 reductions at 128 μM. These findings show that high classifier agreement is not a substitute for genomic evidence and provide an auditable framework for prioritizing both synthetic AMP-like sequences and candidate genome-encoded peptides.

## Introduction

Antimicrobial resistance continues to outpace the development of antibacterial agents with new chemical scaffolds and mechanisms [1–4]. Antimicrobial peptides (AMPs) expand accessible chemical space because activity can arise from compact, compositionally diverse sequences [5–9]. Their translational potential, however, depends on properties that sequence classifiers do not measure, including selectivity, proteolytic stability, pharmacokinetics and activity under defined assay conditions [10–13]. Computational prediction should therefore prioritize experimentally testable molecules rather than imply therapeutic readiness.

Bacterial small proteins offer an attractive but technically challenging starting point. Many are missed or inconsistently annotated by conventional genome-analysis pipelines despite established roles in membrane biology, stress responses, transport and host interaction [14–22]. The same attributes that hinder sORF annotation also complicate peptide discovery: short fragments can occur by chance, contig breaks can truncate coding regions, and homology searches have limited resolving power. A peptide with favourable AMP-like features is therefore not necessarily the product of a complete bacterial gene, and a recurrent coding locus is not necessarily antimicrobial. Conflating these questions leads to avoidable overstatement of evidence.

The scale of public bacterial genome resources now permits systematic discovery, but it also introduces sampling bias because a small number of intensively sequenced species and lineages dominate pooled collections [23,24]. We addressed this by clustering proteins independently within species before defining short representative sequences. We then treated genomic context as an evidence axis distinct from AMP prediction: ORF completeness, proximity to contig termini and recurrence across assemblies were evaluated separately from sequence-level classifier outputs.

This separation is essential because current predictors address different aspects of peptide sequence. AmpScanner v2, Macrel and AMPlify enable high-throughput prioritization from primary sequence, while Macrel also estimates haemolytic propensity [29,31,32]. Their agreement can enrich a candidate set, but it neither validates a gene call nor establishes endogenous expression. Similarly, searches of APD, DRAMP, DBAASP and UniProt describe proximity to existing reference space; they do not establish a new biological function or mechanism [25–28]. We therefore used novelty, physicochemical properties, predicted structures and short molecular-dynamics trajectories as prioritization descriptors, not as functional validation.

Here, we present a genome-context-aware pipeline for bacterial sORF-derived AMP discovery. We construct a species-aware representative resource, apply three-model AMP screening, and audit candidate loci before experimental selection. The workflow deliberately retains two candidate classes: sequence/structure-led peptides, evaluated as synthetic molecules, and genome-supported peptides, which additionally satisfy explicit coding-locus criteria. By connecting these complementary routes to MIC and plate-count MBC experiments, we ask both which sequences are antibacterial under the assay conditions and which have credible genomic support.

## Materials and Methods

### Genome collection, ORF prediction, and within-species clustering

We analysed 649,653 NCBI RefSeq assemblies representing 327 clinically relevant bacterial species. The accession-level manifest, species assignments and species-level counts are provided in Supplementary Table S1. Protein-coding ORFs were predicted for each assembly with Prodigal v2.6.3 in single-genome mode using the standard bacterial genetic code [55]. To limit domination of the candidate space by deeply sampled lineages, predicted proteins were pooled and clustered independently within each species with CD-HIT v4.8.1 at 95% amino-acid identity [56]. The longest member of each cluster was retained as the species-level representative, and representatives 10–100 aa long were retained for screening. Accordingly, all downstream values are representative-record counts, not estimates of the absolute number of short ORFs in a species.

### AMP prediction and consensus tiers

All retained representatives were first evaluated with AmpScanner v2; sequences with an AmpScanner probability >0.5 were then evaluated with Macrel and AMPlify [29,31,32]. We assigned mutually exclusive tiers to preserve the underlying prediction evidence. Tier S comprised sequences positive in all three models and classified as non-haemolytic by Macrel. Tier A comprised either three-model-positive sequences classified as haemolytic by Macrel or AmpScanner–Macrel dual-positive, Macrel-non-haemolytic sequences. Tier B comprised the remaining dual-model-positive sequences. Predictor outputs were treated solely as sequence-level prioritization features; no model score was interpreted as a measurement of antimicrobial activity or gene validity. Full combination counts and tier definitions are provided in Supplementary Table S2.

### Genome-context audit and candidate selection

Candidate occurrence records were audited at the ORF level. An occurrence was classified as edge-associated when the retained project annotation marked the ORF as partial and its coordinate lay within 100 bp of a contig terminus. Complete non-edge support required at least one occurrence that was neither partial nor terminally proximal. The principal genome-supported pool comprised candidates with complete non-edge support in at least two assemblies; the stringent subset comprised candidates observed in at least ten assemblies. Candidate-level counts for complete/non-edge, complete/edge, partial/non-edge and partial/edge occurrences are provided in Supplementary Table S3.

Experimental candidates were selected by two prespecified routes. The sequence/structure-led route started from Tier S and applied novelty, physicochemical and structural-ranking filters. The genome-supported route was drawn from the principal complete non-edge pool. Similarity to known AMP space was assessed against DRAMP, APD, DBAASP and UniProt AMP annotations. N5 novel-family status required both <80% sequence identity and <80% aligned query coverage to the matched reference. This design intentionally distinguishes a sequence selected for synthetic-peptide testing from a sequence additionally supported by a recurrent complete genomic locus.

### Structural and molecular-dynamics prioritization

Selected candidates underwent single-sequence structure prediction with OmegaFold [35]. The resulting models were used as starting conformations for 5-ns all-atom aqueous molecular-dynamics simulations in AMBER14 with TIP3P-FB water at 300 K. Temperature stability, backbone root-mean-square deviation, radius of gyration and potential-energy behaviour were summarized as comparative quality-control descriptors. Because short peptides are conformationally heterogeneous in solution and the trajectories were short, these measurements were used to rank candidates and assess model behaviour; they were not used to assign a membrane-active conformation or an antimicrobial mechanism.

### Peptide synthesis and antimicrobial assays

Twenty peptides were synthesized by Biomatik at stated purity >95%; sequence-level HPLC/MS quality-control records are provided in Supplementary Table S8. MIC screening was performed by broth microdilution in 96-well plates against E. coli ATCC 25922 and S. aureus ATCC 25923. Each peptide was tested at 1, 2, 4, 8, 16, 32, 64 and 128 μM in two technical replicates. Every plate included blank, growth, solvent and meropenem controls. OD600 values were blank-corrected as OD_sample − mean(OD_blank), and the growth reference was mean(OD_growth) − mean(OD_blank). Percent inhibition was calculated as 1 − (corrected sample OD/growth reference); MIC was defined as the lowest tested concentration with ≥90% inhibition in the mean of the two technical replicates. Well-level OD600 data, plate controls and the calling rule are provided in Supplementary Tables S5 and S6.

CAND_04141, CAND_07825 and CAND_04265 were advanced to plate-count MBC assays at five concentrations (8–128 μM). For each peptide–organism–concentration condition, three independent biological replicates were plated at the recorded dilution in 0.1-mL volumes. Colony counts were converted to CFU/mL by dividing the mean colony count by the dilution fraction and plated volume. Log10 reduction was calculated relative to the growth control (6.0 × 10^8 CFU/mL), and MBC was defined as the lowest tested concentration producing a ≥3-log10 reduction. Raw counts, calculated CFU/mL, plate-image traceability and the MBC summary are provided in Supplementary Table S7.

### Statistical reporting

Analyses were descriptive unless otherwise stated. Pearson correlation quantified the association between per-species assembly number and retained sORF representative count. In MBC dose-response displays, points represent biological replicates and lines summarize the arithmetic mean; MBC calls followed the prespecified ≥3-log10 rule rather than a hypothesis test. No inferential comparisons were applied to the two-technical-replicate MIC screen. Exact denominators are reported for every filtering step.

## Results

### A species-aware bacterial sORF resource captures 4.44 million representative short peptides

We collected 649,653 RefSeq bacterial genome assemblies from 327 clinically relevant species. Sampling was highly uneven: the median number of assemblies per species was 78 (interquartile range, 21–402; range, 1–93,149), and the ten most sampled species accounted for 64.9% of all assemblies. ORF prediction with Prodigal [55] yielded approximately 1.19 billion protein-coding predictions, of which approximately 140.7 million encoded 10–100-aa products.

To reduce the influence of repeatedly sampled clonal lineages, proteins were clustered independently within each species at 95% sequence identity using CD-HIT [56], with the longest sequence retained as the cluster representative. This produced 16,731,256 nr95 protein representatives. Length filtering retained 4,442,548 10–100-aa sORF representatives from 320 species (26.6% of all nr95 representatives). The final collection represents a 31.7-fold reduction from the original short-ORF instance count; this reduction reflects both within-species clustering and subsequent representative-length filtering, rather than clustering alone (Figure 1).

**Figure 1.**
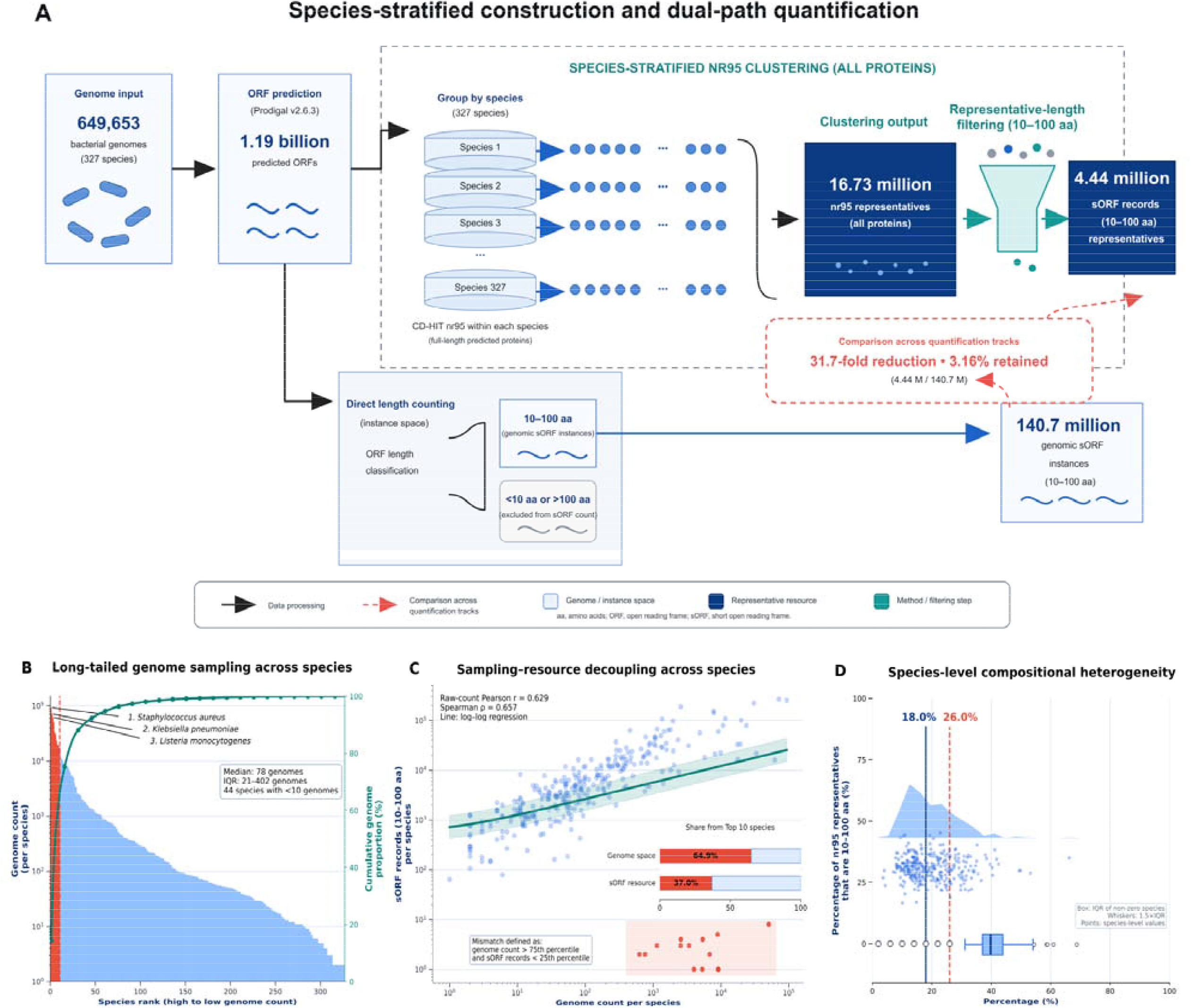
Construction and species-aware compression of a bacterial sORF representative resource. (a) Workflow from genome collection and ORF prediction to within-species nr95 clustering and sORF representative selection. (b –d) Species sampling distribution, relationship between sampling depth and retained sORF diversity, and characteristics of the resulting resource. The final resource contains 4,442,548 10–100-aa representatives from 320 species.

The resource remained sensitive to sampling depth but was not proportional to it. Genome number and final sORF representative number were moderately correlated (Pearson r = 0.629); nevertheless, the ten most sampled species contributed 64.9% of assemblies but only 37.0% of retained sORF representatives. An audit also showed that 15.4% of 10–100-aa cluster members belonged to clusters whose representative exceeded 100 aa and was therefore not retained. Thus, the resource is a conservative discovery set derived from whole-proteome clustering, not a complete census of every short ORF.

This design has two implications for interpretation. First, it limits redundant discovery opportunities in deeply sampled lineages; it is not intended to estimate the absolute abundance of small coding sequences in any species. Second, a species with zero retained short representatives after representative-length filtering should not be interpreted as lacking sORFs. The loss of short members to clusters represented by proteins >100 aa is an explicit consequence of the conservative whole-proteome clustering design. We therefore retain cluster-member traceability and report downstream counts as representative-record counts unless stated otherwise.

### Orthogonal AMP predictors reduce the candidate space to a strict consensus tier

AmpScanner v2 screening (probability >0.5) identified 912,427 canonical-positive records, corresponding to 755,160 exact unique peptide sequences. Macrel and AMPlify were then applied to every AmpScanner-positive sequence. Of these records, 714,718 were positive only in AmpScanner, 184,533 were AmpScanner–AMPlify dual positives, 9,851 were AmpScanner–Macrel dual positives, and 3,325 were supported by all three predictors (Figure 2). AMPlify did not return a prediction for 35,044 sequences containing non-standard amino acids; those records retained only two available model outputs.

**Figure 2.**
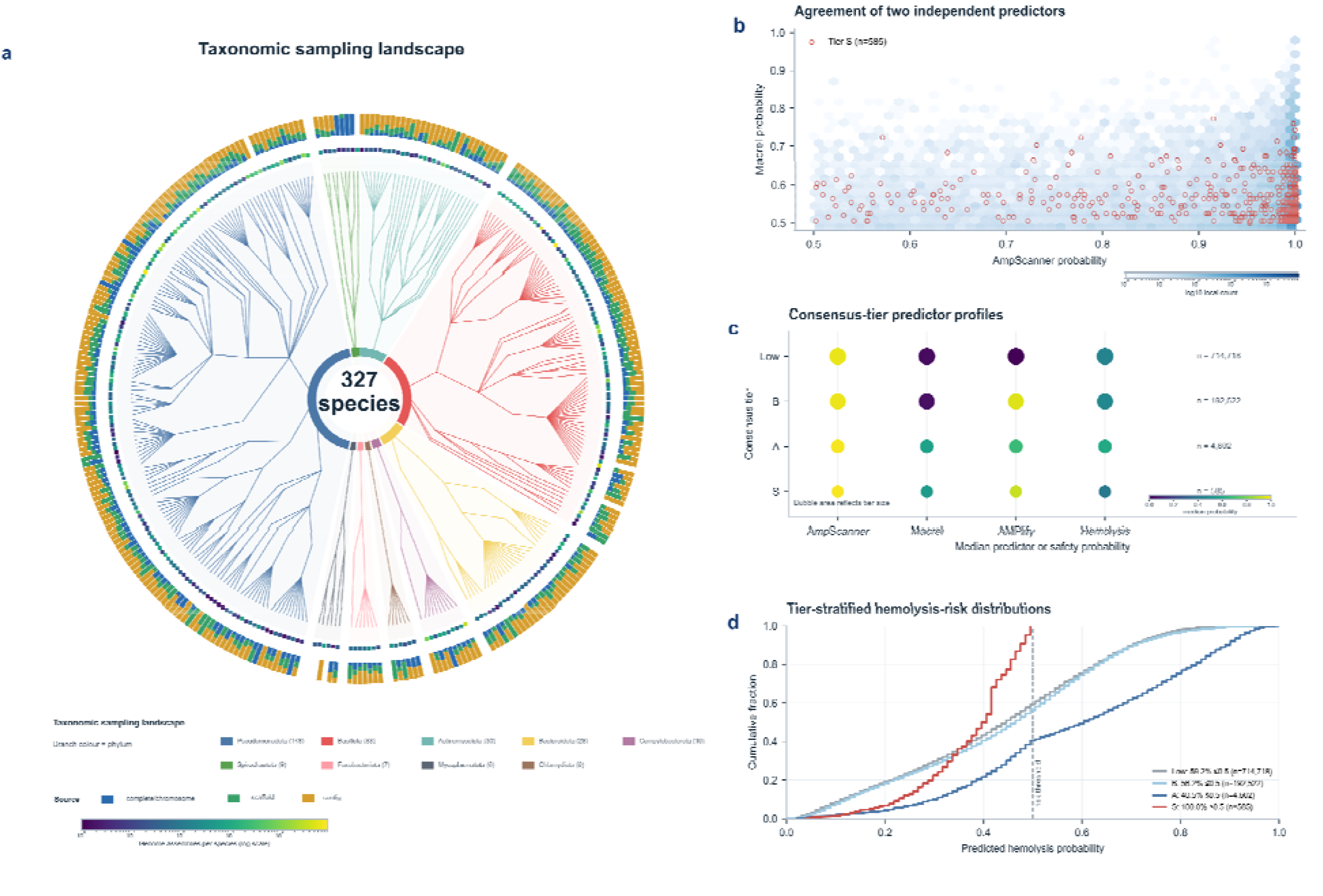
Multi-model consensus stratifies AMP-like sORF candidates. (a) Distribution of predictor-support combinations among AmpScanner-positive records. (b) Tier definitions and sizes. (c–d) Predictor-score and predicted haemolysis landscapes, highlighting Tier S. Model consensus is sequence-level evidence and does not encode genome-context support.

We defined mutually exclusive prediction tiers using model agreement and Macrel haemolysis prediction. Tier S contained 585 records (536 unique sequences) positive in all three models and predicted non-haemolytic. Tier A contained 4,602 records (4,327 unique sequences) that were either three-model positive but predicted haemolytic or AmpScanner–Macrel dual positive and predicted non-haemolytic. Tier B contained 192,522 remaining dual-model-positive records. There was no exact-sequence overlap among tiers. This hierarchy deliberately represents confidence in AMP-like sequence features, rather than proof that a candidate is encoded by a complete bacterial ORF.

The reduction from 912,427 AmpScanner-positive records to 3,325 three-model-positive records illustrates the stringency imposed by orthogonal prediction; it is not a calibrated probability of biological activity. The models are not statistically independent because each learns features related to composition, charge, amphipathicity and sequence patterning. Their agreement is therefore best interpreted as a robust operational filter that enriches a particular AMP-like sequence space, not as three independent experimental confirmations. This distinction is especially important for sORFs, for which the prior probability that a short predicted ORF encodes a stable natural peptide can be low.

### Genome-context auditing reveals a severe edge-partial bias in strict prediction tiers

We next examined genomic occurrence information for 11,918 AmpScanner–Macrel-supported candidates. This traceable set included 536/585 Tier S records, 4,327/4,602 Tier A records and the mapped AmpScanner–Macrel component of Tier B (7,055 records). Among mapped Tier S candidates, 529/536 (98.7%) were supported exclusively by partial ORFs within 100 bp of a contig terminus; none of their 615 mapped instances was complete and non-edge. Only seven Tier S candidates had at least one complete non-edge occurrence. Tier A showed the same, albeit less extreme, pattern: 3,695/4,327 (85.4%) were edge-partial-only and 583 (13.5%) had complete non-edge support. In the mapped Tier B component, 4,108/7,055 (58.2%) were edge-partial-only and 2,792 (39.6%) had complete non-edge support (Supplementary Table S3). These Tier B values apply only to its mapped AmpScanner– Macrel component and are not estimates for the full Tier B population.

Across all 11,918 mapped candidates, 3,382 (28.4%) had at least one complete non-edge genomic occurrence. We designated 1,069 of these candidates as a principal genome-supported pool because they occurred in at least two assemblies; a stringent subset of 251 was observed in at least ten assemblies. Only four Tier S, 185 Tier A, and 880 mapped Tier B candidates comprised the 1,069-candidate pool. Thus, predictor agreement and genomic completeness/reproducibility provided non-redundant evidence, and strict AMP prediction alone was insufficient to nominate well-contextualized ORFs.

The edge-partial enrichment is compatible with several non-exclusive explanations: fragmented assemblies can truncate coding sequences, ORF prediction near contig boundaries is intrinsically uncertain, and short fragments may receive favourable AMP scores because they retain cationic or hydrophobic subsequences. Our analysis cannot distinguish these mechanisms for every record. Its purpose is narrower and operational: the available coordinates do not support classification of edge-partial-only records as complete genomic sORFs. This deliberately conservative audit rule provides a reproducible separation of complete-locus evidence from sequence-only evidence.

### Two complementary prioritization routes assemble an auditable experimental panel

We integrated novelty, physicochemical filtering, structure prediction, aqueous molecular-dynamics (MD) behaviour and genomic support to construct a 20-candidate experimental panel (Figure 3). In the activity/structure-led route, 486 CD-HIT-90 Tier S representatives were evaluated for homology against AMP databases. Of these, 129 (26.5%) were classified as N5 novel-family candidates, defined by <80% identity and <80% coverage to a known AMP. The eight activity/structure-led peptides selected for synthesis were all N5; their best DRAMP identities ranged from 42.9% to 76.5% at 35.0–59.1% coverage.

**Figure 3.**
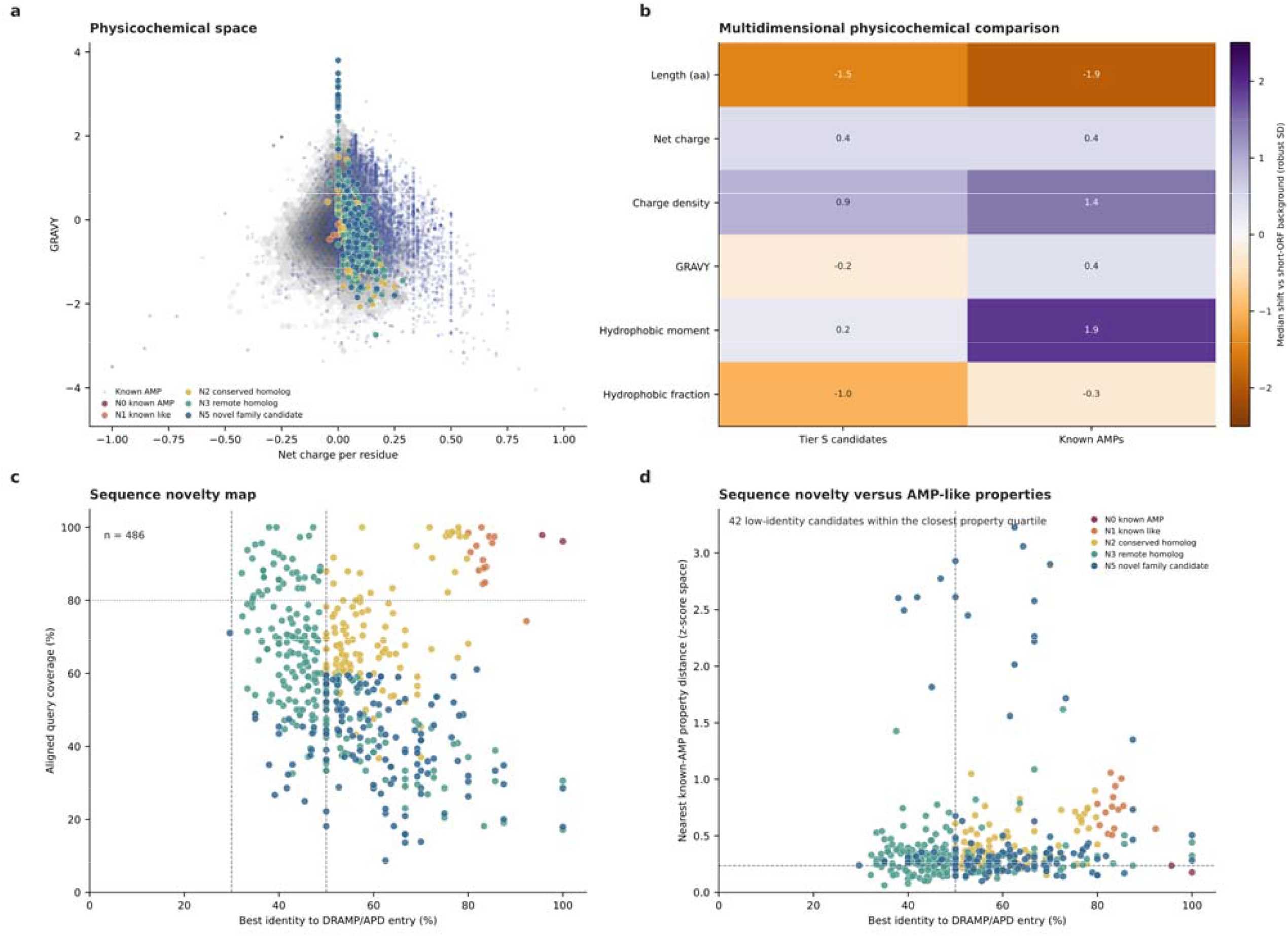
Physicochemical and sequence-novelty landscape of prioritized AMP-like candidates. (a) Net charge per residue and GRAVY distribution, highlighting known AMPs and novelty classes among candidate peptides. (b) Robustly standardized physicochemical features for Tier S candidates and known AMPs. (c) Best identity and aligned query coverage against DRAMP/APD for 486 Tier S representatives; dashed lines mark the classification thresholds shown. (d) Sequence novelty plotted against distance in AMP-like property space; highlighted points identify low-identity candidates within the closest property quartile.

The genome-supported route contributed 12 candidates with complete non-edge support. Eleven had no detectable homology to DRAMP or APD, and one had remote homology (35.2% identity). All 12 had complete non-edge support; 10 occurred in at least two assemblies and nine in at least ten assemblies. The activity/structure-led candidates were retained as peptide-level exploratory candidates because subsequent context auditing showed that all eight were edge-associated partial ORFs. The panel therefore distinguishes a compelling peptide sequence from evidence for a complete genomic coding locus.

Structure prediction and short aqueous MD trajectories were used only as secondary quality-control descriptors. Nineteen Tier S candidates and all 12 genome-supported candidates entered the 5-ns analysis; these descriptors informed prioritization but neither overruled complete-locus evidence nor served as functional evidence. Direct visual assets retained for the eight activity/structure-led experimental peptides with exact-sequence matches to tierS simulation runs are summarized as original-output grids: structural starting conformations (Supplementary Figure S1), Water-MD trajectory plots (Supplementary Figure S2), and membrane-MD endpoint visualizations (Supplementary Figure S3).

### MIC screening identifies active peptides from both selection routes

All 20 synthesized peptides were screened by broth microdilution against E. coli ATCC 25922 and S. aureus ATCC 25923 using a twofold concentration series from 1 to 128 μM and an OD600 readout (Figure 4). Seven candidates had an E. coli MIC ≤16 μM, and eight had an S. aureus MIC ≤4 μM. CAND_04141, a genome-supported candidate from Coxiella burnetii (Tier A; complete non-edge support in 95 assemblies), had the lowest combined MIC: 4 μM against E. coli and 2 μM against S. aureus. CAND_07825, an activity/structure-led Tier S candidate, had MICs of 8 and 2 μM, respectively. CAND_04265, a genome-supported candidate, had MICs of 8 and 4 μM. Thus, both selection routes yielded active peptides, whereas the strongest combined profile came from the genomically recurrent cohort.

**Figure 4.**
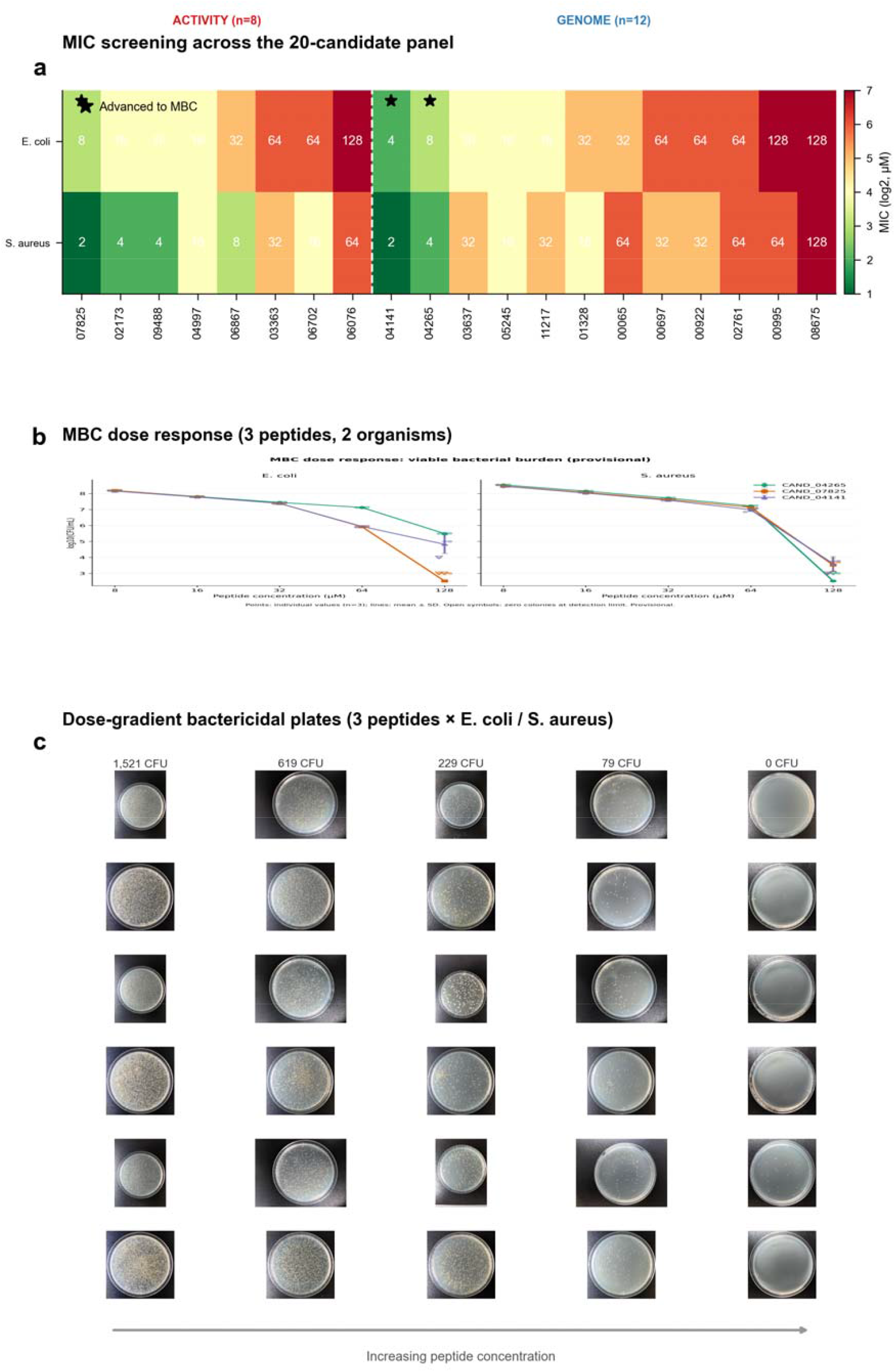
Experimental validation of prioritized pathogen sORF peptides. (a) MIC heatmap for the 20-candidate panel against E. coli ATCC 25922 and S. aureus ATCC 25923. Red and blue bands denote activity/structure-led and genome-supported selection routes, respectively; stars mark peptides advanced to MBC testing. (b) CFU dose response for CAND_04141, CAND_07825 and CAND_04265. Points show independent biological replicates and lines show their arithmetic mean ± SD. (c) Representative plate series for CAND_04141 against E. coli. MBC values were calculated from three independent biological replicates with image-matched colony counts.

Three peptides were advanced to plate-count MBC assessment: CAND_04141 (best combined MIC), CAND_07825 (strong activity in the activity/structure-led cohort) and CAND_04265 (a genome-supported low-micromolar candidate). All three produced concentration-dependent CFU decreases against both organisms, and all six peptide–organism combinations reached a ≥3-log10 reduction at 128 μM (Figure 4). The MBC endpoint was 128 μM for every combination (MBC/MIC ratios, 16–64). MBC data are based on three independent biological replicates with image-matched colony counts.

The MIC patterns further indicate that activity was not captured by a single computational selection variable. CAND_04141 was selected from the genome-supported route despite Tier A classification, whereas CAND_07825 originated from the strict Tier S activity/structure-led route. The two organisms did not respond uniformly: MIC differences for a given peptide ranged from two-to sixteenfold, consistent with organism-specific interactions between peptide properties and envelope architecture. This study was not designed to infer mechanism from these patterns, but they justify prioritizing the three peptides for follow-up testing under standardized peptide-assay conditions, including low-binding plasticware and appropriate medium controls [39–42].

## Discussion

The central finding is not merely the recovery of antibacterial sequences, but the extent of disagreement between sequence-model consensus and genomic coding evidence. Nearly all mapped Tier S candidates were represented exclusively by partial ORFs near contig termini. This distinction matters because small proteins are already difficult to annotate and validate in bacterial genomes [14–22]. If classifier agreement is treated as evidence of gene identity, edge-associated fragments can be inadvertently promoted from sequence hypotheses to putative endogenous products. The observed pattern is compatible with fragmented assemblies, uncertain gene calling at termini and enrichment of short cationic or hydrophobic fragments by sequence classifiers. It is therefore a methodological finding with broad relevance to genome-scale AMP mining, not a peculiarity of one prediction tool.

The context audit does not diminish the value of sequence-led discovery. CAND_07825 arose from the sequence/structure-led route and showed low-micromolar MICs, establishing antibacterial activity of the synthesized sequence under the conditions tested. It does not, however, establish that the associated partial ORF is expressed as the same mature peptide in its source organism. Conversely, CAND_04141 combined the strongest two-strain activity profile with a complete non-edge locus recurring in 95 assemblies. This convergence of chemical activity and locus-level evidence makes CAND_04141 the most defensible current candidate for follow-up as a putative genome-encoded AMP. The two routes are therefore complementary: one explores chemically promising sequence space, whereas the other prioritizes candidates for which a genomic-origin claim is supportable.

These results support a practical reporting standard for computational peptide discovery. Sequence prediction, locus confidence and experimental function should be retained as separate, inspectable fields rather than merged into a single confidence score. Predictor ensembles can be valuable because their decision boundaries differ [29,31,32], but consensus does not make the models independent or substitute for genomic context. Likewise, low similarity to AMP databases describes current reference-space coverage, not proof of a new biological mechanism [25–28]. Maintaining these distinctions improves interpretability and enables comparisons across studies without conflating synthetic activity with endogenous expression.

The experimental data establish a clear next-stage prioritization, but do not constitute a complete preclinical package. The MIC screen identifies an initial activity pattern across a Gram-negative and a Gram-positive reference strain, and plate-count MBC assays demonstrate concentration-dependent killing for the three advanced peptides across three independent biological replicates. Apparent potency in peptide assays can nevertheless vary with inoculum, medium, plate material, endpoint and host-cell interactions [12,39–42]. Moreover, short aqueous MD trajectories cannot establish a membrane-bound structure or mechanism. The appropriate interpretation is therefore that this study identifies experimentally supported antibacterial hits and a locus-aware prioritization framework, not therapeutic readiness.

The most informative next experiments are therefore decision-focused. Repeating MIC and MBC assays with independently prepared cultures and peptide dilutions will test reproducibility. Haemolysis, mammalian-cell toxicity and serum/protease stability will determine whether antibacterial activity is accompanied by an acceptable initial safety window, while time-kill, membrane-permeabilization and resistance-selection experiments will examine mechanism and liability [10–13,43–46]. For genome-supported candidates, ribosome profiling, targeted proteomics or reporter assays should determine whether the predicted loci are transcribed and translated. Together, these data would resolve the two remaining biological questions: whether the active synthetic sequences are tractable leads, and whether the strongest genome-supported candidates are naturally produced by their source organisms.

## Conclusions

Genome-context-aware discovery separates two claims that are often merged in sORF mining: that a short sequence is antibacterial when synthesized and that it is encoded by a credible bacterial locus. Applied to 649,653 assemblies, the framework exposed a pronounced edge-partial bias among strict sequence-prediction hits while yielding a 20-peptide experimental panel with multiple low-micromolar MICs. CAND_04141 is the leading genome-supported candidate. This approach is readily transferable to other bacterial genome-mining studies and provides a transparent basis for the expression, safety and mechanistic experiments required before translational interpretation.

## Supporting information

Pathogen_sORF_AMP_Supplementary_Information

Pathogen_sORF_AMP_Supplementary_Code

Pathogen_sORF_AMP_Supplementary_Tables

## Data Availability

The processed source data supporting the conclusions, including Supplementary Tables S1–S8 and figure source data, are supplied with this article and its Supplementary Materials. Genome assemblies were retrieved from NCBI RefSeq. Additional raw files and candidate-ID mappings are available from the corresponding author upon reasonable request.

## Code Availability

Custom scripts used for data assembly, MBC calculation and figure generation are supplied as Supplementary Code. The full genome-scale workflow additionally requires the original RefSeq collection and the external predictor installations described in the Methods.

## Ethics, Competing Interests, and Author Contributions

### Ethics approval and consent to participate

Not applicable.

### Author contributions

Qingxiu Li: Conceptualization, methodology, data curation, formal analysis, visualization, investigation, and writing—original draft. Zhenjun Li: Supervision, project administration, interpretation of results, writing—review and editing, and correspondence.

### Funding

This research received no specific grant from any funding agency in the public, commercial, or not-for-profit sectors.

