## Supplementary material for "Genome-context-aware discovery of antibacterial peptides from bacterial small open reading frames": Pathogen_sORF_AMP_Supplementary_Information

Supplementary Figures S1–S3 and Supplementary Tables S1–S8

### Supplementary Figure S1 | Original structural starting conformations for eight mapped activity/structure-led experimental peptides.

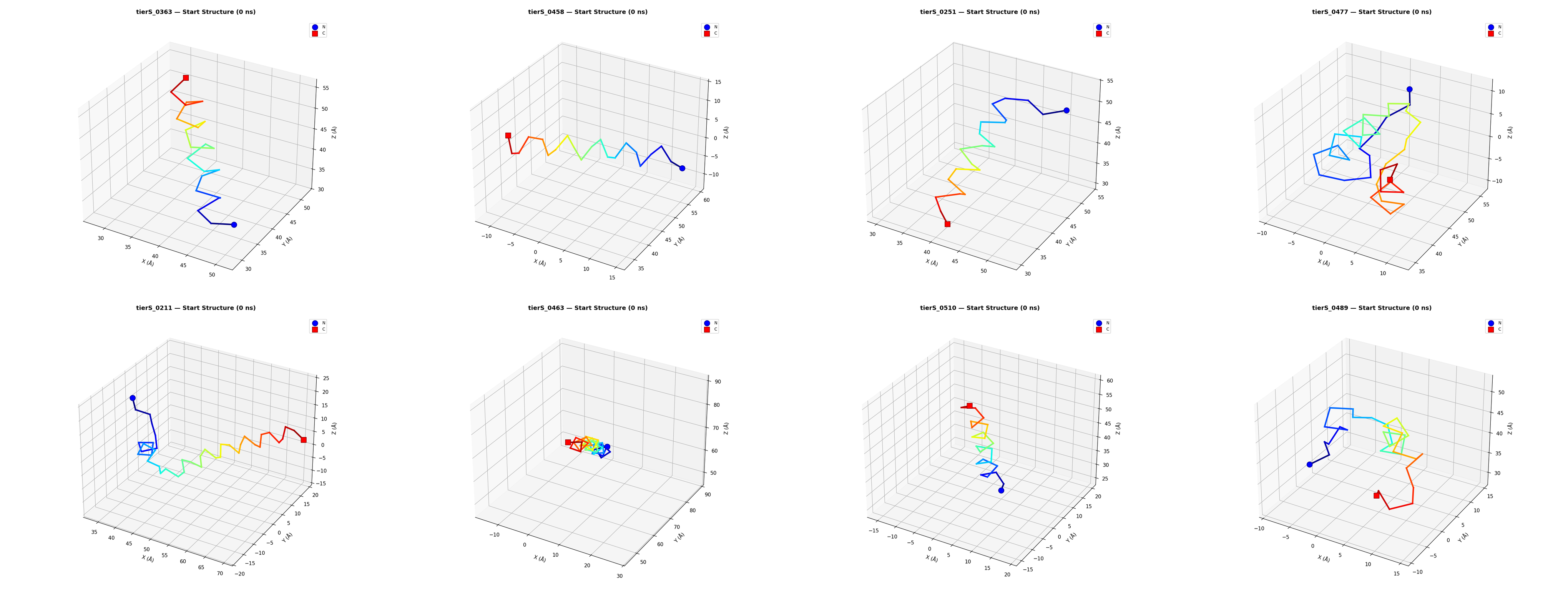

### Supplementary Figure S2 | Original 5-ns Water-MD trajectory summaries for the same eight peptides.

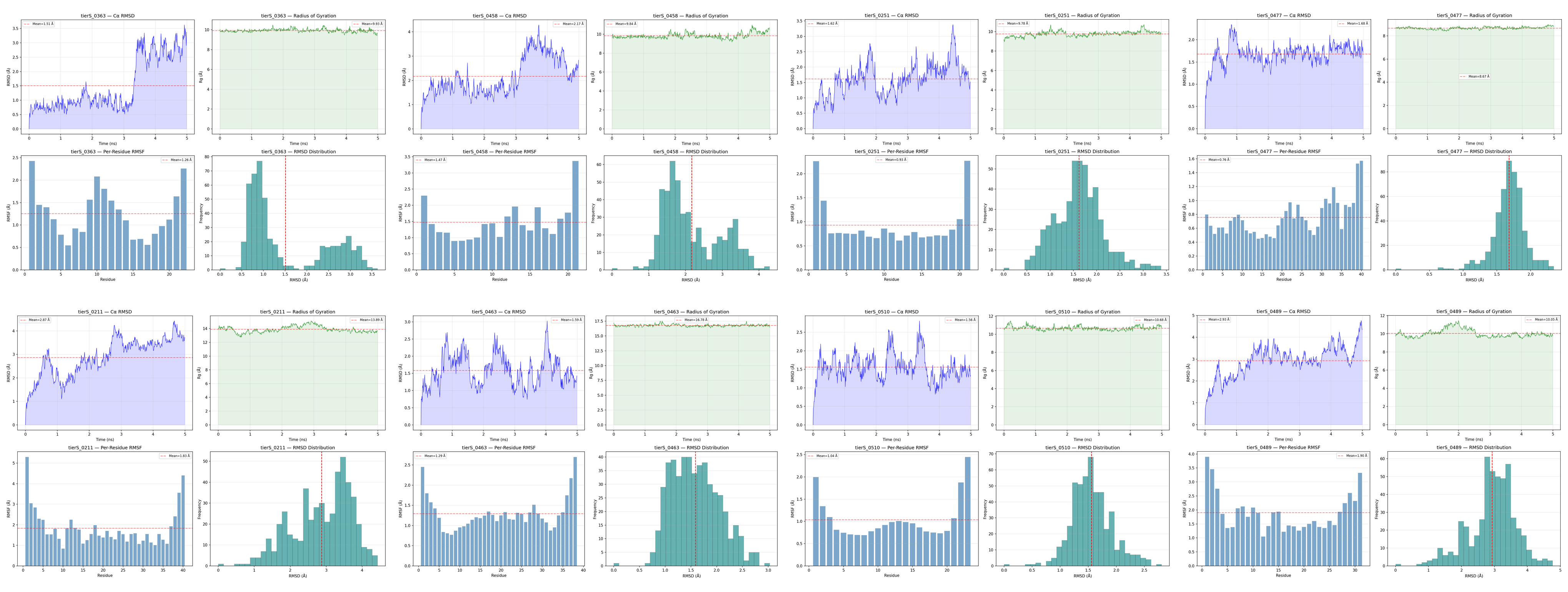

### Supplementary Figure S3 | Original membrane-MD endpoint visualizations for the same eight peptides.

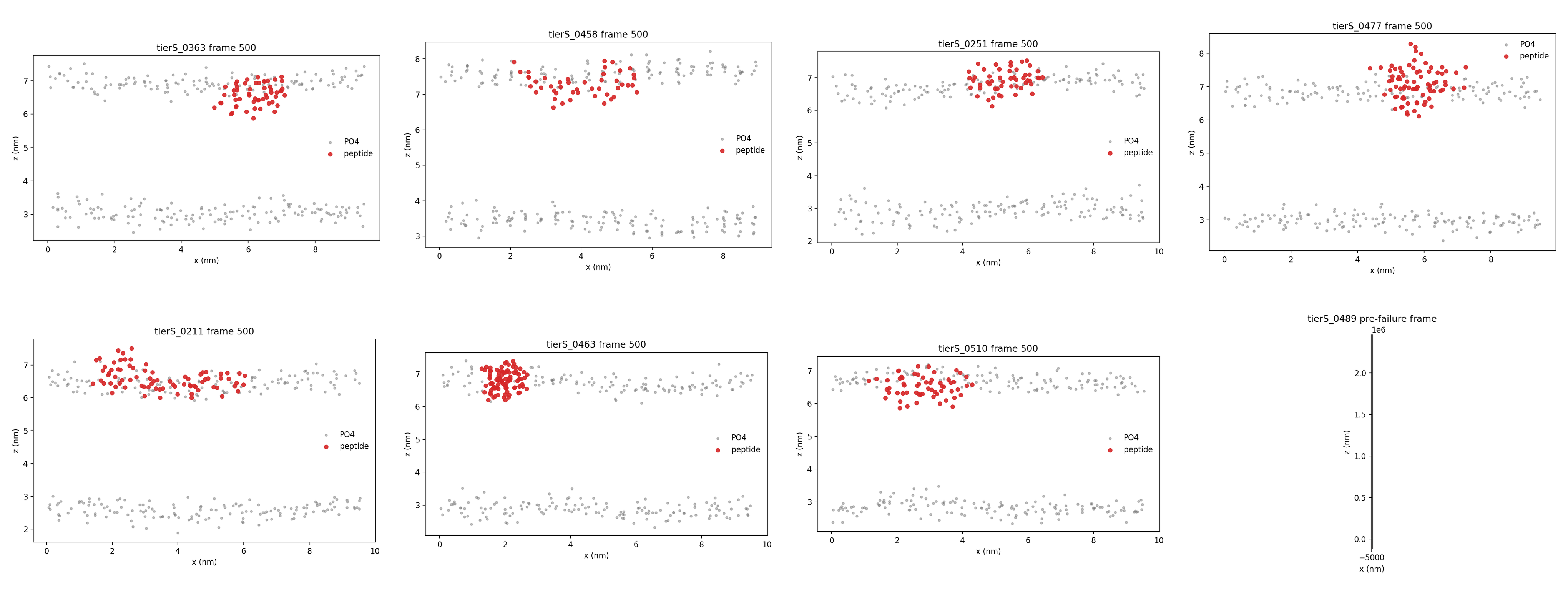

### Supplementary Tables

Supplementary Tables S1–S8 are supplied as the accompanying archive Pathogen_sORF_AMP_Supplementary_Tables.zip. The archive includes editable Excel workbooks and tab-separated source files. Table S7 contains the three-replicate MBC data: Rep1, Rep2, and Rep3 colony counts are all image-matched and verified.
