## Supplementary material for "Genome-context-aware discovery of antibacterial peptides from bacterial small open reading frames": Pathogen_sORF_AMP_Supplementary_Code: README.md

# Supplementary Code

This archive contains the project scripts used to assemble the submission tables, figures, MBC calculations, and this preprint package. The scripts retain their original project-relative input paths. Processed source tables are supplied separately in `Pathogen_sORF_AMP_Supplementary_Tables.zip`.

## Environment

Scripts were run with Python 3 and use standard-library modules plus pandas, numpy, matplotlib, openpyxl, and python-docx where applicable. The large genome source collection and external predictor installations are not redistributed in this archive; accession-level summaries and processed results are provided in the Supplementary Tables.

## Reproducibility scope

The archive reproduces data assembly, calculation and figure-generation steps from retained project inputs. Rerunning the full genome-scale workflow requires the original RefSeq collection and the installed predictor software.
