## Supplementary material for "Genome-context-aware discovery of antibacterial peptides from bacterial small open reading frames": Pathogen_sORF_AMP_Supplementary_Code: build_biorxiv_submission_package.py

```
#!/usr/bin/env python3
"""Create a bioRxiv-oriented submission package from the audited manuscript assets."""

from pathlib import Path
import hashlib
import os
import re
import shutil
import subprocess
from zipfile import ZIP_DEFLATED, ZipFile

from docx import Document
from docx.enum.text import WD_ALIGN_PARAGRAPH
from docx.shared import Inches, Pt, RGBColor

ROOT = Path(__file__).resolve().parents[1]
MANUSCRIPT = ROOT / "manuscript_submission" / "SCI_MANUSCRIPT_FULL_DRAFT.md"
SUPP = ROOT / "figures_generated" / "supplementary"
TABLES = SUPP / "tables"
MAIN = ROOT / "figures_generated" / "main_figures_nc_v2"
MAIN_V2 = ROOT / "figures_generated" / "main_figures_v2"
MD_SUMMARY = ROOT / "figures_generated" / "supplementary" / "original_md_summary"
OUT = Path(os.environ.get("BIORXIV_PACKAGE_OUT", ROOT / "manuscript_submission" / "bioRxiv_submission_package"))
CODE_FILES = [
    ROOT / "scripts" / "build_submission_supplementary_tables.py",
    ROOT / "scripts" / "plot_supplementary_figures.py",
    ROOT / "scripts" / "build_final_supplementary_composites.py",
    ROOT / "scripts" / "confirm_mbc_s7_metadata.py",
    ROOT / "scripts" / "build_biorxiv_submission_package.py",
    ROOT / "scripts" / "build_original_md_summary_figures.py",
    ROOT / "section_3_5" / "04_analysis" / "build_mbc_optimized_traceable.py",
    ROOT / "section_3_5" / "04_analysis" / "build_mbc_standardized_table.py",
    ROOT / "section_3_5" / "04_analysis" / "reconcile_new_mic_files_to_old_source.py",
]

TITLE = "Genome-context-aware discovery of antibacterial peptides from bacterial small open reading frames"
AUTHORS = "Qingxiu Li¹, Zhenjun Li¹*"
AFFILIATION = "¹Institute of Infectious Diseases, Chinese Center for Disease Control and Prevention, 155 Changbai Road, Changping District, Beijing 102206, China"
CORRESPONDING = "*Correspondence: Zhenjun Li"


def clean_md(text):
    text = text.strip()
    text = text.replace("**", "").replace("`", "")
    text = re.sub(r"\*([^*]+)\*", r"\1", text)
    return text


def configure(doc):
    section = doc.sections[0]
    section.top_margin = Inches(0.8)
    section.bottom_margin = Inches(0.8)
    section.left_margin = Inches(0.85)
    section.right_margin = Inches(0.85)
    normal = doc.styles["Normal"]
    normal.font.name = "Times New Roman"
    normal.font.size = Pt(10.5)
    normal.paragraph_format.space_after = Pt(6)
    normal.paragraph_format.line_spacing = 1.1
    for name, size in [("Title", 17), ("Heading 1", 13), ("Heading 2", 11.5)]:
        style = doc.styles[name]
        style.font.name = "Arial"
        style.font.size = Pt(size)
        style.font.bold = True


def add_para(doc, text, bold=False, center=False):
    p = doc.add_paragraph()
    p.alignment = WD_ALIGN_PARAGRAPH.CENTER if center else WD_ALIGN_PARAGRAPH.LEFT
    run = p.add_run(clean_md(text))
    run.bold = bold
    return p


def manuscript_docx(path):
    doc = Document()
    configure(doc)
    p = doc.add_paragraph(style="Title")
    p.alignment = WD_ALIGN_PARAGRAPH.CENTER
    p.add_run(TITLE)
    add_para(doc, AUTHORS, center=True)
    add_para(doc, AFFILIATION, center=True)
    add_para(doc, CORRESPONDING, center=True)
    p = add_para(doc, "bioRxiv preprint | This manuscript has not been peer reviewed.", center=True)
    p.runs[0].font.color.rgb = RGBColor(120, 0, 0)
    doc.add_page_break()

    text = MANUSCRIPT.read_text(encoding="utf-8")
    body = "## Abstract" + text.split("## Abstract", 1)[1].split("## Submission-critical items", 1)[0]
    # Render the conventional SCI research-article sequence even while the
    # working Markdown is being edited in-place by the project team.
    parts = re.split(r"(?=^## )", body, flags=re.MULTILINE)
    by_heading = {part.splitlines()[0][3:].strip(): part for part in parts if part.startswith("## ")}
    ordered = [
        "Abstract", "Introduction", "Materials and Methods", "Results", "Discussion", "Conclusions",
        "Data Availability", "Code Availability", "Ethics, Competing Interests, and Author Contributions",
        "Figure Legends", "Tables", "References",
    ]
    body = "\n\n".join(by_heading[name] for name in ordered if name in by_heading)
    for raw in body.splitlines():
        line = raw.strip()
        if not line:
            continue
        if line.startswith("### "):
            doc.add_heading(clean_md(line[4:]), level=2)
        elif line.startswith("## "):
            doc.add_heading(clean_md(line[3:]), level=1)
        elif re.match(r"^\d+\. ", line):
            doc.add_paragraph(clean_md(re.sub(r"^\d+\. ", "", line)), style="List Number")
        elif line.startswith("- "):
            doc.add_paragraph(clean_md(line[2:]), style="List Bullet")
        elif line.startswith("|") or line.startswith("---"):
            # The manuscript has no data tables in the main text; their source files are supplied separately.
            continue
        else:
            add_para(doc, line)

    doc.add_page_break()
    doc.add_heading("Main Figures", level=1)
    figures = [
        ("Figure 1", MAIN / "Figure1_source.png"),
        ("Figure 2", MAIN / "Figure_2_multimodel_consensus.png"),
        ("Figure 3", MAIN / "Main_Fig3_physicochemical_novelty.png"),
        ("Figure 4", MAIN_V2 / "Main_v2_Fig4_experimental_validation.png"),
    ]
    for label, fig in figures:
        if not fig.exists() and label == "Figure 4":
            fig = MAIN / "Main_Fig4_experimental_validation.png"
        doc.add_heading(label, level=2)
        doc.add_picture(str(fig), width=Inches(6.7))
        doc.add_page_break()
    doc.save(path)


def supplementary_docx(path):
    doc = Document()
    configure(doc)
    p = doc.add_paragraph(style="Title")
    p.alignment = WD_ALIGN_PARAGRAPH.CENTER
    p.add_run("Supplementary Information")
    add_para(doc, TITLE, bold=True, center=True)
    add_para(doc, "Supplementary Figures S1–S3 and Supplementary Tables S1–S8", center=True)
    doc.add_page_break()
    figures = [
        ("Supplementary Figure S1 | Original structural starting conformations for eight mapped activity/structure-led experimental peptides.", MD_SUMMARY / "Supplementary_FigS1_original_structures_8_candidates.png"),
        ("Supplementary Figure S2 | Original 5-ns Water-MD trajectory summaries for the same eight peptides.", MD_SUMMARY / "Supplementary_FigS2_original_water_MD_8_candidates.png"),
        ("Supplementary Figure S3 | Original membrane-MD endpoint visualizations for the same eight peptides.", MD_SUMMARY / "Supplementary_FigS3_original_membrane_MD_8_candidates.png"),
    ]
    for title, fig in figures:
        doc.add_heading(title, level=1)
        doc.add_picture(str(fig), width=Inches(6.7))
        doc.add_page_break()
    doc.add_heading("Supplementary Tables", level=1)
    add_para(doc, "Supplementary Tables S1–S8 are supplied as the accompanying archive `Pathogen_sORF_AMP_Supplementary_Tables.zip`. The archive includes editable Excel workbooks and tab-separated source files. Table S7 contains the three-replicate MBC data: Rep1, Rep2, and Rep3 colony counts are all image-matched and verified.")
    doc.save(path)


def supplementary_code_zip(path):
    readme = """# Supplementary Code

This archive contains the project scripts used to assemble the submission tables, figures, MBC calculations, and this preprint package. The scripts retain their original project-relative input paths. Processed source tables are supplied separately in `Pathogen_sORF_AMP_Supplementary_Tables.zip`.

## Environment

Scripts were run with Python 3 and use standard-library modules plus pandas, numpy, matplotlib, openpyxl, and python-docx where applicable. The large genome source collection and external predictor installations are not redistributed in this archive; accession-level summaries and processed results are provided in the Supplementary Tables.

## Reproducibility scope

The archive reproduces data assembly, calculation and figure-generation steps from retained project inputs. Rerunning the full genome-scale workflow requires the original RefSeq collection and the installed predictor software.
"""
    with ZipFile(path, "w", ZIP_DEFLATED) as archive:
        archive.writestr("README.md", readme)
        for file in CODE_FILES:
            if file.exists():
                archive.write(file, arcname=f"scripts/{file.name}")


def convert_pdf(docx_path):
    profile = Path("/tmp") / "codex_biorxiv_lo_profile"
    profile.mkdir(exist_ok=True)
    subprocess.run([
        "libreoffice", f"-env:UserInstallation=file://{profile}", "--headless",
        "--convert-to", "pdf", "--outdir", str(docx_path.parent), str(docx_path),
    ], check=True, stdout=subprocess.DEVNULL)


def sha256(path):
    h = hashlib.sha256()
    with path.open("rb") as handle:
        for block in iter(lambda: handle.read(1 << 20), b""):
            h.update(block)
    return h.hexdigest()


def write_text(path, text):
    path.write_text(text, encoding="utf-8")


def main():
    if OUT.exists():
        shutil.rmtree(OUT)
    (OUT / "01_manuscript").mkdir(parents=True)
    (OUT / "02_supplementary").mkdir()
    (OUT / "03_metadata_and_declarations").mkdir()
    subprocess.run(["python3", str(ROOT / "scripts" / "build_original_md_summary_figures.py")], check=True)

    manuscript = OUT / "01_manuscript" / "Pathogen_sORF_AMP_bioRxiv_preprint.docx"
    supplement = OUT / "02_supplementary" / "Pathogen_sORF_AMP_Supplementary_Information.docx"
    manuscript_docx(manuscript)
    supplementary_docx(supplement)
    convert_pdf(manuscript)
    convert_pdf(supplement)

    table_zip = OUT / "02_supplementary" / "Pathogen_sORF_AMP_Supplementary_Tables.zip"
    with ZipFile(table_zip, "w", ZIP_DEFLATED) as archive:
        for file in sorted(TABLES.iterdir()):
            if file.is_file():
                archive.write(file, arcname=file.name)
    supplementary_code_zip(OUT / "02_supplementary" / "Pathogen_sORF_AMP_Supplementary_Code.zip")
    shutil.copy2(MD_SUMMARY / "MD_asset_manifest.tsv", OUT / "02_supplementary" / "MD_asset_manifest.tsv")

    metadata = """# bioRxiv submission metadata — complete before upload

| Field | Proposed / required value |
|---|---|
| Title | Genome-context-aware discovery of antibacterial peptides from bacterial small open reading frames |
| Article type | New results / Research article |
| Primary category | Microbiology |
| Suggested keywords | small open reading frame; antimicrobial peptide; bacterial genomes; genome context; machine learning; peptide discovery |
| Abstract | Copy the finalized Abstract from the manuscript PDF; do not alter wording after PDF generation. |
| Authors | Qingxiu Li; Zhenjun Li |
| Affiliations | Institute of Infectious Diseases, Chinese Center for Disease Control and Prevention, Beijing, China |
| Corresponding author | Zhenjun Li; |
| ORCID | Not provided; add if available. |
| Funding | No external funding: “This research received no specific grant from any funding agency in the public, commercial, or not-for-profit sectors.” |
| Competing interests | The authors declare no competing interests. |
| License | Recommended: CC-BY 4.0, subject to all authors’ and funders’ agreement. |
| Prior dissemination | None. |
| Data / code link | Processed data are supplied as Supplementary Materials. Reproducible assembly, calculation and figure-generation scripts are supplied as `Pathogen_sORF_AMP_Supplementary_Code.zip`; additional raw files are available from the corresponding author upon reasonable request. |
"""
    write_text(OUT / "03_metadata_and_declarations" / "bioRxiv_submission_metadata_TEMPLATE.md", metadata)

    author_template = "author_order\tfull_name\taffiliation_number\taffiliation_text\tORCID\tcorresponding_author\temail\n1\tQingxiu Li\t1\tInstitute of Infectious Diseases, Chinese Center for Disease Control and Prevention, 155 Changbai Road, Changping District, Beijing 102206, China\t[not provided]\tNO\\n2\tZhenjun Li\t1\tInstitute of Infectious Diseases, Chinese Center for Disease Control and Prevention, 155 Changbai Road, Changping District, Beijing 102206, China\t[not provided]\tYES\\n"
    write_text(OUT / "03_metadata_and_declarations" / "author_affiliation_TEMPLATE.tsv", author_template)

    declarations = """# Statements to confirm before bioRxiv upload

## Competing interests
The authors declare no competing interests.

## Funding
This research received no specific grant from any funding agency in the public, commercial, or not-for-profit sectors.

## Ethics
Not applicable: this study used public bacterial genome assemblies and in-vitro bacterial assays and did not involve human participants, identifiable human data, or vertebrate animals. **Confirm institutional requirements.**

## Data and code availability
The preprint package contains the processed tables, figure files and reproducible assembly, calculation and figure-generation scripts. Additional raw files and candidate-ID mappings are available from the corresponding author upon reasonable request. A public repository DOI may be added in a later version if one becomes available.

## MBC data
Supplementary Table S7 reports the three-replicate MBC data: Rep1, Rep2, and Rep3 are independent biological replicates with image-matched colony counts. MBC endpoints use the prespecified ≥3-log10 reduction criterion.
"""
    write_text(OUT / "03_metadata_and_declarations" / "submission_declarations_TEMPLATE.md", declarations)

    files = [p for p in OUT.rglob("*") if p.is_file()]
    manifest_rows = ["relative_path\tbytes\tsha256\tpurpose"]
    for f in sorted(files):
        rel = f.relative_to(OUT)
        purpose = "submission file" if f.suffix.lower() in {".pdf", ".zip"} else "metadata/template"
        manifest_rows.append(f"{rel}\t{f.stat().st_size}\t{sha256(f)}\t{purpose}")
    write_text(OUT / "03_metadata_and_declarations" / "file_manifest.tsv", "\n".join(manifest_rows) + "\n")

    readme = """# bioRxiv submission package

## Upload order

1. Upload `01_manuscript/Pathogen_sORF_AMP_bioRxiv_preprint.pdf` as the main manuscript.
2. Upload `02_supplementary/Pathogen_sORF_AMP_Supplementary_Information.pdf` as supplementary information.
3. Upload `02_supplementary/Pathogen_sORF_AMP_Supplementary_Tables.zip` as supplementary data.
4. Upload `02_supplementary/Pathogen_sORF_AMP_Supplementary_Code.zip` as supplementary code.
5. Enter metadata using `03_metadata_and_declarations/bioRxiv_submission_metadata_TEMPLATE.md`.
6. Keep the Word files locally; they are editable sources and are not required as the display version.

## Blocking items before clicking submit

- Confirm the CRediT author-contribution and competing-interest statements.
- Confirm that the manuscript’s data/code-availability wording accurately reflects what can be supplied on reasonable request.
- Confirm all authors approve the exact PDF, author order, category, and selected license.
- Confirm the three-replicate MBC data are approved for the preprint version.

## Contents

- Main manuscript: formatted PDF and editable DOCX, with Figures 1–4 embedded.
- Supplementary information: formatted PDF and editable DOCX, with three grids assembled solely from original structure/MD PNGs.
- Supplementary data: Tables S1–S8 in XLSX/TSV format, packaged as ZIP.
- `MD_asset_manifest.tsv`: exact candidate-to-run mappings for the original structure/MD grids.
- Supplementary code: assembly, calculation and figure-generation scripts, packaged as ZIP.
- Metadata/declaration templates and SHA-256 file manifest.

This package is ready for author completion and final approval, not for unattended automated submission.
"""
    write_text(OUT / "README_UPLOAD_ORDER.md", readme)
    print(f"[OK] bioRxiv package created: {OUT}")


if __name__ == "__main__":
    main()
```
