## Supplementary material for "Genome-context-aware discovery of antibacterial peptides from bacterial small open reading frames": Pathogen_sORF_AMP_Supplementary_Code: build_final_supplementary_composites.py

```
#!/usr/bin/env python3
"""Build the two merged supplementary figures without modifying source figures."""
from pathlib import Path
import matplotlib
matplotlib.use("Agg")
import matplotlib.pyplot as plt
import matplotlib.image as mpimg
import numpy as np
import pandas as pd

ROOT = Path("/media/qxl/EAGET/Pathogen_data/Pathogen_AMP_Mining/01_sorf_amp")
SC = ROOT / "02_screening_consensus"
OUT = ROOT / "00_PROJECT_MASTER/results_section_audit/figures_generated/supplementary/supplementary_final"
OUT.mkdir(parents=True, exist_ok=True)
INK, BLUE, RED, ORANGE, GREY, LIGHT_BLUE, GREEN = "#20242A", "#2C7BB6", "#D7191C", "#FDAE61", "#969696", "#ABD9E9", "#1A9641"
plt.rcParams.update({"font.family": "DejaVu Sans", "font.size": 7, "axes.linewidth": .6,
                     "axes.spines.top": False, "axes.spines.right": False, "svg.fonttype": "none"})

def panel(ax, path, label):
    img = mpimg.imread(path)
    # Source exports often contain broad white margins. Crop these mechanically
    # so labels and marks remain readable in the assembled supplementary figure.
    rgb = img[..., :3]
    mask = np.any(rgb < .985, axis=2)
    yy, xx = np.where(mask)
    if len(xx):
        pad = max(8, int(min(img.shape[:2]) * .015))
        img = img[max(0, yy.min()-pad):min(img.shape[0], yy.max()+pad),
                  max(0, xx.min()-pad):min(img.shape[1], xx.max()+pad)]
    ax.imshow(img)
    ax.axis("off")
    ax.text(-.025, 1.025, label, transform=ax.transAxes, fontsize=9, fontweight="bold", va="bottom")

def save(fig, stem):
    for ext in ("png", "pdf", "svg"):
        fig.savefig(OUT / f"{stem}.{ext}", dpi=600 if ext == "png" else None, facecolor="white")
    plt.close(fig)

def fig_s2():
    ra = ROOT / "00_PROJECT_MASTER/results_section_audit"
    gen, s33 = ra / "figures_generated", ra / "section_3_3/03_main_figures"
    fig = plt.figure(figsize=(7.1, 8.7), constrained_layout=True)
    grid = fig.add_gridspec(3, 2)
    sources = [
        (grid[0, 0], s33 / "Figure_3a_audit_funnel.png", "a", "Audit funnel"),
        (grid[0, 1], s33 / "Figure_3b_genomic_support_by_tier.png", "b", "Genomic support by tier"),
        (grid[1, 0], gen / "Fig3C_genomic_evidence_composition.png", "c", "Evidence composition"),
        (grid[1, 1], gen / "Fig3D_gene_neighborhood.png", "d", "Gene neighborhood"),
        (grid[2, :], gen / "Fig3E_sequence_conservation_logo.png", "e", "Sequence conservation"),
    ]
    for slot, image, label, title in sources:
        ax = fig.add_subplot(slot); panel(ax, image, label); ax.set_title(title, loc="left", fontsize=7)
    fig.suptitle("Supplementary Fig. S2 | Genomic-context and edge-bias quality control",
                 fontsize=10, fontweight="bold", color=INK)
    save(fig, "Supplementary_FigS2_genome_context_QC")

def fig_s3():
    d04 = SC / "04_sequence_cluster_robustness/figures"
    d07 = SC / "07_pangenome_analysis/figures"
    d07r = SC / "07_pangenome_analysis_refined/figures"
    fig = plt.figure(figsize=(7.1, 8.6), constrained_layout=True)
    grid = fig.add_gridspec(3, 2, height_ratios=[.82, 1, 1])
    panels = [
        (grid[0, 0], d04 / "fig1_coverage_sensitivity.png", "a"),
        (grid[0, 1], d04 / "fig2_length_analysis.png", "b"),
        (grid[1, 0], d07 / "fig1_gram_comparison.png", "c"),
        (grid[1, 1], d07 / "fig2_eskape_comparison.png", "d"),
        (grid[2, 0], d07r / "fig1_cliffs_delta_gram.png", "e"),
        (grid[2, 1], d07r / "fig2_zero_inflation.png", "f"),
    ]
    for slot, image, label in panels:
        panel(fig.add_subplot(slot), image, label)
    fig.suptitle("Supplementary Fig. S3 | Robustness to sequence clustering and pangenome refinement",
                 fontsize=10, fontweight="bold", color=INK)
    save(fig, "Supplementary_FigS3_clustering_pangenome_robustness")

def fig_s5():
    matrix = pd.read_csv(SC / "05_exact_presence_absence/full_revalidation/near_ubiquitous_full_evidence_matrix.tsv", sep="\t")
    audit = pd.read_csv(SC / "validation/outputs/cross_genus_identical_nt_audit.tsv", sep="\t")
    fig = plt.figure(figsize=(7.1, 6.0), constrained_layout=True)
    grid = fig.add_gridspec(2, 3, height_ratios=[.82, 1])
    top = fig.add_subplot(grid[0, :]); top.axis("off")
    cols = ["candidate_id", "evidence_completeness", "experimental_priority"]
    table = top.table(cellText=matrix[cols].values.tolist(), colLabels=["Candidate", "Evidence completeness", "Priority"],
                      cellLoc="left", colLoc="left", loc="center")
    table.auto_set_font_size(False); table.set_fontsize(7); table.scale(1, 1.32)
    for (row, col), cell in table.get_celld().items():
        cell.set_linewidth(.35); cell.set_edgecolor(GREY)
        if row == 0:
            cell.set_facecolor(LIGHT_BLUE); cell.get_text().set_weight("bold")
    top.text(-.025, 1.01, "a", transform=top.transAxes, fontsize=9, fontweight="bold")
    top.set_title("Near-ubiquitous candidates are retained as provisional exceptions", loc="left", fontsize=8, pad=4)

    ax = fig.add_subplot(grid[1, 0]); vc = audit["classification"].value_counts()
    ax.bar(range(len(vc)), vc.values, color=[BLUE, GREY, ORANGE][:len(vc)], linewidth=0)
    ax.set(xticks=range(len(vc)), xticklabels=[x.replace("_", "\n") for x in vc.index], ylabel="Candidates")
    ax.tick_params(axis="x", labelsize=5.7)
    for i, value in enumerate(vc.values): ax.text(i, value, str(value), ha="center", va="bottom", fontsize=6)
    ax.text(-.16, 1.04, "b", transform=ax.transAxes, fontsize=9, fontweight="bold")
    ax.set_title("Audit classification", loc="left", fontsize=8)

    ax = fig.add_subplot(grid[1, 1]); ax.hist(audit["n_species"], bins=np.arange(2, 8) - .5, color=ORANGE, edgecolor="white", linewidth=.3)
    ax.set(xlabel="Species containing candidate", ylabel="Candidates")
    ax.text(-.16, 1.04, "c", transform=ax.transAxes, fontsize=9, fontweight="bold")
    ax.set_title("Cross-species extent", loc="left", fontsize=8)

    ax = fig.add_subplot(grid[1, 2]); values = [audit["nt_identical"].astype(bool).sum(), audit["plasmid_signal"].astype(bool).sum()]
    ax.bar(["nt-identical", "plasmid\nsignal"], values, color=[GREEN, RED], linewidth=0)
    ax.set_ylabel("Candidates")
    for i, value in enumerate(values): ax.text(i, value, str(value), ha="center", va="bottom", fontsize=6)
    ax.text(-.16, 1.04, "d", transform=ax.transAxes, fontsize=9, fontweight="bold")
    ax.set_title("Potential artifact signals", loc="left", fontsize=8)
    fig.suptitle("Supplementary Fig. S5 | Revalidation and artifact audit of exceptional candidates",
                 fontsize=10, fontweight="bold", color=INK)
    save(fig, "Supplementary_FigS5_exception_candidate_audit")

def fig_s6():
    water = pd.read_csv(SC / "05_water_md/water_md_summary.tsv", sep="\t")
    water["candidate_id"] = water.query_id.astype(str).str.split("|").str[0]
    vis = SC / "05_water_md/analysis_visualization"
    memvis = SC / "06_membrane_md/visualizations"
    colors = {"PASS": "#2A9D8F", "BORDERLINE": "#E9B949", "FAIL": "#C65D5D"}
    groups = {key: water.loc[water.qc_class.eq(key), "candidate_id"].tolist() for key in colors}
    fig = plt.figure(figsize=(18, 12.6), constrained_layout=True)
    grid = fig.add_gridspec(4, 8)
    fig.suptitle("Supplementary Fig. S5 | Structure-prediction and molecular-dynamics quality-control gallery",
                 fontsize=13, fontweight="bold", color=INK)
    rows = [("a", "Water-MD PASS: start–end structural superpositions", "PASS", "superposition"),
            ("b", "Membrane-MD final frames for water-MD PASS candidates", "PASS", "membrane"),
            ("c", "Water-MD BORDERLINE: start–end structural superpositions", "BORDERLINE", "superposition"),
            ("d", "Water-MD FAIL: original structural inspection renders", "FAIL", "cartoon")]
    for row, (letter, title, status, mode) in enumerate(rows):
        fig.text(.01, .948 - row*.24, f"{letter}  {title}", fontsize=10, fontweight="bold")
        for col, candidate in enumerate(groups[status]):
            if col >= 8: break
            ax = fig.add_subplot(grid[row, col])
            if mode == "superposition": path = vis / status / candidate / f"{candidate}_superposition.png"
            elif mode == "cartoon": path = vis / status / candidate / f"{candidate}_overall_cartoon.png"
            elif candidate == "tierS_0489": path = memvis / "tierS_0489_failed_prefailure_xz.png"
            else: path = memvis / f"{candidate}_frame_0500_xz.png"
            if path.exists(): ax.imshow(mpimg.imread(path))
            else: ax.text(.5, .5, "image unavailable", ha="center", va="center", fontsize=6)
            ax.set_title(candidate.replace("tierS_", ""), fontsize=6.8, color=colors[status], fontweight="bold", pad=2); ax.axis("off")
    fig.text(.5, .012, "Images are original pipeline outputs; PASS/BORDERLINE/FAIL denote water-MD QC classes.", ha="center", fontsize=7.2, color=GREY)
    save(fig, "Supplementary_FigS5_structure_MD_gallery")

if __name__ == "__main__":
    # S2 remains a renamed source export until its archived component panels
    # are restored; the other figures are rebuilt from available source data.
    fig_s3(); fig_s5(); fig_s6()
```
