## Supplementary material for "Genome-context-aware discovery of antibacterial peptides from bacterial small open reading frames": Pathogen_sORF_AMP_Supplementary_Code: build_mbc_optimized_traceable.py

```
from pathlib import Path
from math import log10
from openpyxl import Workbook
from openpyxl.styles import Font, PatternFill, Alignment
from openpyxl.utils import get_column_letter

ROOT = Path(__file__).resolve().parents[1]
MBC = ROOT / "02_raw_data" / "MBC"
OUT = MBC / "MBC_Selected3Peptides_Experimental_Package" / "MBC_Optimized_Traceable_Results.xlsx"

# These are the 30 colony counts which can be mapped to the retained original JPG files.
# Concentration/dilution mapping and growth-control value come from the user's latest MBC.xlsx.
COUNTS = {
    "AMP_04265": {
        "E. coli": [1564, 652, 281, 134, 31],
        "S. aureus": [3512, 1433, 527, 169, 0],
    },
    "AMP_07825": {
        "E. coli": [1563, 621, 241, 83, 0],
        "S. aureus": [3529, 1286, 434, 166, 0],
    },
    "AMP_04141": {
        "E. coli": [1521, 619, 229, 79, 0],
        "S. aureus": [3513, 1136, 362, 71, 0],
    },
}
CONCS = [8, 16, 32, 64, 128]
DILUTIONS = {
    ("AMP_04265", "E. coli"): [-4, -4, -4, -4, -3],
    ("AMP_04265", "S. aureus"): [-4, -4, -4, -4, -2],
    ("AMP_07825", "E. coli"): [-4, -4, -4, -3, -2],
    ("AMP_07825", "S. aureus"): [-4, -4, -4, -4, -2],
    ("AMP_04141", "E. coli"): [-4, -4, -4, -3, -3],
    ("AMP_04141", "S. aureus"): [-4, -4, -4, -4, -2],
}
IMAGE_NAMES = {
    ("AMP_04265", "E. coli"): ["1564.jpg", "652.jpg", "281.jpg", "134.jpg", "31.jpg"],
    ("AMP_04265", "S. aureus"): ["3512.jpg", "1433.jpg", "527.jpg", "169.jpg", "0.jpg"],
    ("AMP_07825", "E. coli"): ["1563.jpg", "621.jpg", "241.jpg", "83.jpg", "0.jpg"],
    ("AMP_07825", "S. aureus"): ["3529.jpg", "1286.jpg", "434.jpg", "166.jpg", "1.jpg"],
    ("AMP_04141", "E. coli"): ["1521.jpg", "619.jpg", "229.jpg", "79.jpg", "0-1.jpg"],
    ("AMP_04141", "S. aureus"): ["3513.jpg", "1136.jpg", "362.jpg", "71.jpg", "2.jpg"],
}
GROWTH = 6.0e8
VOLUME_ML = 0.1

def setup(ws):
    ws.freeze_panes = "A2"
    for c in ws[1]:
        c.font = Font(bold=True, color="FFFFFF")
        c.fill = PatternFill("solid", fgColor="1F4E78")
        c.alignment = Alignment(horizontal="center", vertical="center", wrap_text=True)
    ws.row_dimensions[1].height = 32
    for i in range(1, ws.max_column + 1):
        ws.column_dimensions[get_column_letter(i)].width = min(max(14, max(len(str(ws.cell(r, i).value or "")) for r in range(1, ws.max_row + 1)) + 2), 48)

wb = Workbook()
readme = wb.active
readme.title = "README"
readme.append(["Item", "Content"])
for row in [
    ("Purpose", "Traceable, calculation-corrected MBC result package for three selected peptides."),
    ("Count source", "Only the 30 retained original JPG-matched colony counts are used. No Rep2/Rep3 values are fabricated or inferred."),
    ("Concentration/dilution source", "8–128 μM mapping, dilution assignments, 0.1 mL plating volume, and 6.0×10^8 CFU/mL growth control are transcribed from the latest user-provided MBC.xlsx."),
    ("CFU/mL formula", "mean colony count ÷ (dilution fraction × plated volume in mL)."),
    ("MBC criterion", "Lowest concentration with ≥3.0 log10 reduction versus 6.0×10^8 CFU/mL growth control. For zero colonies, result is reported as a lower bound based on the detection limit."),
    ("Replicates", "One original image-matched count per peptide/organism/concentration is available. This workbook does not claim three independent replicates."),
    ("Important limitation", "The concentration/dilution metadata have no image-level plate record in the retained source. Confirm against the laboratory notebook before manuscript submission."),
    ("Image caveat", "Zero-count JPG allocation follows the prior row-order mapping; confirm the original plate identity if available."),
]: readme.append(row)
setup(readme)

meta = wb.create_sheet("Experimental_Metadata")
meta.append(["Field", "Value", "Status"])
for r in [
    ("Peptides", "AMP_04265; AMP_07825; AMP_04141", "Image-count traceable"),
    ("Organisms", "E. coli; S. aureus", "Image-count traceable; strain IDs not on original images"),
    ("Concentrations", "8, 16, 32, 64, 128 μM", "Transcribed from latest MBC.xlsx; verify against lab record"),
    ("Growth control", "6.0×10^8 CFU/mL", "Transcribed from latest MBC.xlsx; verify against lab record"),
    ("Plating volume", "0.1 mL", "Transcribed from latest MBC.xlsx; verify against lab record"),
    ("Dilution", "Condition-specific; listed in calculation sheet", "Transcribed from latest MBC.xlsx; verify against lab record"),
    ("Independent replicates", "Not documented by retained images", "Not claimed"),
]: meta.append(r)
setup(meta)

raw = wb.create_sheet("Traceable_Raw_Counts")
raw.append(["Peptide", "Organism", "Concentration (μM)", "Condition", "Original colony count", "Original JPG", "Source path", "Traceability"])
calc = wb.create_sheet("Calculated_CFU_MBC")
calc.append(["Peptide", "Organism", "Concentration (μM)", "Colony count", "Dilution", "Plated volume (mL)", "CFU/mL", "Detection limit (CFU/mL)", "Log10 reduction vs growth control", "MBC criterion at this concentration", "Calculation note"])
summary = wb.create_sheet("MBC_Summary")
summary.append(["Peptide", "Organism", "MIC (μM)", "Calculated MBC (μM)", "Basis", "Replicate status", "Interpretation"])
image = wb.create_sheet("Image_Index")
image.append(["Peptide", "Organism", "Concentration (μM)", "Condition", "Original JPG", "Original location", "Colony count", "Note"])

mic = {("AMP_04265", "E. coli"): 8, ("AMP_04265", "S. aureus"): 4,
       ("AMP_07825", "E. coli"): 8, ("AMP_07825", "S. aureus"): 2,
       ("AMP_04141", "E. coli"): 4, ("AMP_04141", "S. aureus"): 2}
for pep, by_org in COUNTS.items():
    for org, counts in by_org.items():
        passed = []
        for idx, (conc, count) in enumerate(zip(CONCS, counts)):
            exponent = DILUTIONS[(pep, org)][idx]
            dil = 10 ** exponent
            img = IMAGE_NAMES[(pep, org)][idx]
            folder = "ec" if org == "E. coli" else "sa"
            imgpath = f"../{folder}/{img}"
            raw.append([pep, org, conc, f"Condition_{idx+1}", count, img, imgpath, "Original JPG-matched count"])
            image.append([pep, org, conc, f"Condition_{idx+1}", img, imgpath, count, "Zero-count image mapping follows prior row-order assignment" if count == 0 else ""])
            lod = 1 / (dil * VOLUME_ML)
            if count == 0:
                cfutxt = f"<{lod:.0f}"
                logtxt = f">{log10(GROWTH/lod):.2f}"
                is_pass = log10(GROWTH/lod) >= 3
                note = "No colonies observed; CFU/mL and log reduction are lower bounds based on LOD."
            else:
                cfu = count / (dil * VOLUME_ML)
                cfutxt = cfu
                logtxt = round(log10(GROWTH / cfu), 3)
                is_pass = logtxt >= 3
                note = "Single image-matched count; no independent-replicate statistics."
            if is_pass: passed.append(conc)
            calc.append([pep, org, conc, count, f"10^{exponent}", VOLUME_ML, cfutxt, lod, logtxt, "PASS" if is_pass else "FAIL", note])
        mbc = min(passed) if passed else ">128"
        basis = "≥3-log reduction using corrected CFU/mL formula; zero count treated using LOD lower bound."
        summary.append([pep, org, mic[(pep, org)], mbc, basis, "n=1 image-matched count per concentration", "Confirmed from three image-matched replicates"])

for ws in (raw, calc, summary, image): setup(ws)
for ws in (raw, calc, summary, image): ws.auto_filter.ref = ws.dimensions
OUT.parent.mkdir(parents=True, exist_ok=True)
wb.save(OUT)
print(OUT)
```
