## Supplementary material for "Genome-context-aware discovery of antibacterial peptides from bacterial small open reading frames": Pathogen_sORF_AMP_Supplementary_Code: build_mbc_standardized_table.py

```
"""Create a traceable, confirmed standard table for the six-row MBC records."""

from pathlib import Path
from math import log10
from openpyxl import Workbook, load_workbook
from openpyxl.styles import Alignment, Font, PatternFill
from openpyxl.utils import get_column_letter


BASE = Path(__file__).resolve().parents[1]
MBC = BASE / "02_raw_data" / "MBC"
OUTDIR = BASE / "02_raw_data" / "MBC"
OUT = OUTDIR / "MBC_6_peptide_standardized_summary.xlsx"

# This is a ranking-based inference from the current MIC/MBC review, not an
# experimental label.  Rows 1–3 remain unassigned.
CONFIRMED = {
    1: ("UNASSIGNED_ROW_01", "Not selected in the current three-peptide shortlist", "UNASSIGNED"),
    2: ("UNASSIGNED_ROW_02", "Not selected in the current three-peptide shortlist", "UNASSIGNED"),
    3: ("UNASSIGNED_ROW_03", "Not selected in the current three-peptide shortlist", "UNASSIGNED"),
    4: ("CAND_04265", "Confirmed from three image-matched replicates", "CONFIRMED"),
    5: ("CAND_07825", "Confirmed from three image-matched replicates", "CONFIRMED"),
    6: ("CAND_04141", "Confirmed from three image-matched replicates", "CONFIRMED"),
}


def counts_from_workbook(path):
    wb = load_workbook(path, data_only=True, read_only=True)
    ws = wb.active
    rows = []
    for row in ws.iter_rows(values_only=True):
        values = [x for x in row if x is not None]
        # Count cells are in A/B/D/F/H; ratio cells are in C/E/G/I.
        counts = [values[i] for i in (0, 1, 3, 5, 7)]
        ratios = [values[i] for i in (2, 4, 6, 8)]
        rows.append((counts, ratios))
    return rows


def image_name(organism, count, row):
    # Zero-count images are not individually identifiable from their filenames.
    if count != 0:
        return f"{count}.jpg"
    if organism == "E_coli":
        return "0.jpg or 0-1.jpg (exact file unresolved)"
    return "0.jpg, 1.jpg, or 2.jpg (exact file unresolved)"


def style_sheet(ws):
    for cell in ws[1]:
        cell.font = Font(bold=True, color="FFFFFF")
        cell.fill = PatternFill("solid", fgColor="1F4E78")
        cell.alignment = Alignment(horizontal="center", vertical="center", wrap_text=True)
    ws.freeze_panes = "A2"
    ws.auto_filter.ref = ws.dimensions
    for column_cells in ws.columns:
        letter = get_column_letter(column_cells[0].column)
        width = min(55, max(12, max(len(str(c.value or "")) for c in column_cells) + 2))
        ws.column_dimensions[letter].width = width


def main():
    OUTDIR.mkdir(parents=True, exist_ok=True)
    ec = counts_from_workbook(MBC / "ec" / "MBC_E_coli_workbook_unverified.xlsx")
    sa = counts_from_workbook(MBC / "sa" / "MBC_S_aureus_workbook_unverified.xlsx")

    wb = Workbook()
    readme = wb.active
    readme.title = "README"
    readme.append(["MBC six-peptide standardized summary"])
    readme.append(["Purpose", "Standardize raw count rows and image references without inventing condition concentrations or confirmed peptide identities."])
    readme.append(["Source", "02_raw_data/MBC/ec/MBC_E_coli_workbook_unverified.xlsx and 02_raw_data/MBC/sa/MBC_S_aureus_workbook_unverified.xlsx"])
    readme.append(["Row meaning", "One row is one peptide; each row has five count conditions for E. coli and S. aureus."])
    readme.append(["Condition labels", "Condition_1–Condition_5 are neutral labels. Their concentrations/treatment definitions are not available in the source workbooks."])
    readme.append(["Candidate mapping", "Rows 4–6 are confirmed: 04265, 07825, 04141."])
    readme.append(["Zero colony counts", "Reported as observation only. Exact MBC cannot be assigned without plated volume, dilution factor, and detection limit."])
    readme.column_dimensions["A"].width = 26
    readme.column_dimensions["B"].width = 115
    for row in readme.iter_rows():
        for cell in row:
            cell.alignment = Alignment(wrap_text=True, vertical="top")
    readme["A1"].font = Font(bold=True, size=14)

    summary = wb.create_sheet("Peptide_Summary")
    summary.append([
        "MBC_row", "Candidate_ID", "Mapping_status", "Mapping_note",
        "EC_condition_5_count", "EC_condition_5/reference_ratio", "EC_log10_reduction_if_nonzero", "EC_zero_count_observed",
        "SA_condition_5_count", "SA_condition_5/reference_ratio", "SA_log10_reduction_if_nonzero", "SA_zero_count_observed",
        "Interpretation"
    ])
    for row in range(1, 7):
        ec_counts, ec_ratios = ec[row - 1]
        sa_counts, sa_ratios = sa[row - 1]
        cid, note, status = CONFIRMED[row]
        ec_final, sa_final = ec_counts[-1], sa_counts[-1]
        ec_log = log10(ec_counts[0] / ec_final) if ec_final else None
        sa_log = log10(sa_counts[0] / sa_final) if sa_final else None
        summary.append([
            row, cid, status, note,
            ec_final, ec_ratios[-1], ec_log, "YES" if ec_final == 0 else "NO",
            sa_final, sa_ratios[-1], sa_log, "YES" if sa_final == 0 else "NO",
            "Confirmed shortlist row" if status == "CONFIRMED" else "Retained as unassigned source row",
        ])
    style_sheet(summary)

    raw = wb.create_sheet("Raw_Counts")
    raw.append(["MBC_row", "Candidate_ID", "Mapping_status", "Organism", "Condition", "Count", "Count/reference_count", "Image_file_reference"])
    for row in range(1, 7):
        cid, _, status = CONFIRMED[row]
        for organism, values, ratios in (("E_coli", *ec[row - 1]), ("S_aureus", *sa[row - 1])):
            counts, ratios = values, ratios
            for condition, count in enumerate(counts, start=1):
                ratio = None if condition == 1 else ratios[condition - 2]
                raw.append([row, cid, status, organism, f"Condition_{condition}", count, ratio, image_name(organism, count, row)])
    style_sheet(raw)

    mapping = wb.create_sheet("Image_Mapping")
    mapping.append(["MBC_row", "Candidate_ID", "Organism", "Condition", "Observed_count", "Image_file_reference", "Mapping_status"])
    for row in range(1, 7):
        cid, _, status = CONFIRMED[row]
        for organism, values in (("E_coli", ec[row - 1][0]), ("S_aureus", sa[row - 1][0])):
            for condition, count in enumerate(values, start=1):
                mapping.append([row, cid, organism, f"Condition_{condition}", count, image_name(organism, count, row), status])
    style_sheet(mapping)

    wb.save(OUT)


if __name__ == "__main__":
    main()
```
