## Supplementary material for "Genome-context-aware discovery of antibacterial peptides from bacterial small open reading frames": Pathogen_sORF_AMP_Supplementary_Code: build_original_md_summary_figures.py

```
#!/usr/bin/env python3
"""Assemble three supplementary grids from retained, unmodified MD visual outputs.

The grids contain the eight activity/structure-led experimental peptides that
map by exact sequence to existing tierS structure/MD outputs. No panel is
redrawn: ImageMagick only lays out the original PNG files in a 4 × 2 grid.
"""

from pathlib import Path
import subprocess

ROOT = Path(__file__).resolve().parents[1]
SCREEN = ROOT.parents[1] / "02_screening_consensus"
WATER = SCREEN / "05_water_md" / "analysis_visualization" / "PASS"
MEMBRANE = SCREEN / "06_membrane_md" / "visualizations"
OUT = ROOT / "figures_generated" / "supplementary" / "original_md_summary"

# Exact-sequence mapping retained by the project: experimental candidate -> tierS run.
RUNS = [
    ("CAND_07825", "tierS_0363"),
    ("CAND_02173", "tierS_0458"),
    ("CAND_06867", "tierS_0251"),
    ("CAND_09488", "tierS_0477"),
    ("CAND_04997", "tierS_0211"),
    ("CAND_03363", "tierS_0463"),
    ("CAND_06702", "tierS_0510"),
    ("CAND_06076", "tierS_0489"),
]


def require(paths):
    missing = [str(path) for path in paths if not path.exists()]
    if missing:
        raise FileNotFoundError("Missing retained source assets:\n" + "\n".join(missing))


def montage(inputs, output):
    require(inputs)
    subprocess.run([
        "montage", *map(str, inputs), "-tile", "4x2", "-geometry", "1000x750+10+10",
        "-background", "white", str(output),
    ], check=True)


def main():
    OUT.mkdir(parents=True, exist_ok=True)
    structural = [WATER / run / f"{run}_start_0ns.png" for _, run in RUNS]
    water_md = [WATER / run / f"{run}_analysis_plots.png" for _, run in RUNS]
    membrane_md = [
        MEMBRANE / (f"{run}_failed_prefailure_xz.png" if run == "tierS_0489" else f"{run}_frame_0500_xz.png")
        for _, run in RUNS
    ]

    montage(structural, OUT / "Supplementary_FigS1_original_structures_8_candidates.png")
    montage(water_md, OUT / "Supplementary_FigS2_original_water_MD_8_candidates.png")
    montage(membrane_md, OUT / "Supplementary_FigS3_original_membrane_MD_8_candidates.png")

    manifest = OUT / "MD_asset_manifest.tsv"
    lines = ["candidate_id\ttierS_run\tstructure_png\twater_md_png\tmembrane_md_png\twater_md_gif"]
    for candidate, run in RUNS:
        membrane_name = f"{run}_failed_prefailure_xz.png" if run == "tierS_0489" else f"{run}_frame_0500_xz.png"
        lines.append("\t".join([
            candidate, run,
            str(WATER / run / f"{run}_start_0ns.png"),
            str(WATER / run / f"{run}_analysis_plots.png"),
            str(MEMBRANE / membrane_name),
            str(WATER / run / f"{run}_5ns_animation.gif"),
        ]))
    manifest.write_text("\n".join(lines) + "\n", encoding="utf-8")

    print(f"[OK] original MD summary assets created: {OUT}")


if __name__ == "__main__":
    main()
```
