## Supplementary material for "Genome-context-aware discovery of antibacterial peptides from bacterial small open reading frames": Pathogen_sORF_AMP_Supplementary_Code: build_submission_supplementary_tables.py

```
#!/usr/bin/env python3
"""Build traceable Supplementary Tables S1-S4 for the manuscript.

Every value is copied or deterministically joined from audited project tables.
No experimental value is imputed.  Run from any directory.
"""

from pathlib import Path
import pandas as pd
from openpyxl.styles import Alignment, Font, PatternFill
from openpyxl.utils import get_column_letter

ROOT = Path(__file__).resolve().parents[1]
OUT = ROOT / "figures_generated" / "supplementary" / "tables"


def read_tsv(relative):
    return pd.read_csv(ROOT / relative, sep="\t", dtype=str, keep_default_na=False)


def style_workbook(path):
    """Apply simple, editable submission formatting after pandas export."""
    from openpyxl import load_workbook

    wb = load_workbook(path)
    for ws in wb.worksheets:
        ws.freeze_panes = "A2"
        ws.auto_filter.ref = ws.dimensions
        for cell in ws[1]:
            cell.font = Font(bold=True, color="FFFFFF")
            cell.fill = PatternFill("solid", fgColor="1F4E78")
            cell.alignment = Alignment(horizontal="center", vertical="center", wrap_text=True)
        ws.row_dimensions[1].height = 34
        for col_idx, column_cells in enumerate(ws.iter_cols(), 1):
            values = [str(c.value or "") for c in column_cells[:200]]
            width = min(max(11, max(map(len, values), default=11) + 2), 45)
            ws.column_dimensions[get_column_letter(col_idx)].width = width
            for cell in column_cells[1:]:
                cell.alignment = Alignment(vertical="top", wrap_text=True)
    wb.save(path)


def write_book(stem, sheets):
    xlsx = OUT / f"{stem}.xlsx"
    with pd.ExcelWriter(xlsx, engine="openpyxl") as writer:
        for sheet_name, frame in sheets.items():
            frame.to_excel(writer, index=False, sheet_name=sheet_name)
    style_workbook(xlsx)


def main():
    OUT.mkdir(parents=True, exist_ok=True)

    # Table S1: complete per-species resource table and the corresponding summary statistics.
    s1 = read_tsv("section_3_1/figures/Figure1_sampling_aware/data/F1_species_level_analysis.tsv")
    s1_summary = read_tsv("section_3_1/figures/Figure1_sampling_aware/data/F1_statistics.tsv")
    s1.to_csv(OUT / "Supplementary_Table_S1_species_sORF_resource_statistics.tsv", sep="\t", index=False)
    s1_summary.to_csv(OUT / "Supplementary_Table_S1_summary_statistics.tsv", sep="\t", index=False)
    write_book("Supplementary_Table_S1_species_sORF_resource_statistics", {
        "S1_species_resource": s1,
        "S1_summary_statistics": s1_summary,
    })

    # Table S2: exact model-support combinations and the audited screen flow.
    s2_support = read_tsv("section_3_2/data/SUPPLEMENTARY_TABLE_3_2_MODEL_SUPPORT_COMBINATIONS.tsv")
    s2_flow = read_tsv("section_3_2/data/SUPPLEMENTARY_TABLE_3_2_SCREENING_FLOW.tsv")
    s2_support.to_csv(OUT / "Supplementary_Table_S2_model_support_combinations.tsv", sep="\t", index=False)
    s2_flow.to_csv(OUT / "Supplementary_Table_S2_screening_flow.tsv", sep="\t", index=False)
    write_book("Supplementary_Table_S2_model_consensus", {
        "S2_model_support": s2_support,
        "S2_screening_flow": s2_flow,
    })

    # Table S3: candidate-level genomic occurrence audit; fields are preserved without collapsing.
    s3 = read_tsv("section_3_3/01_master/candidate_genomic_support_classification.tsv")
    numeric = s3[["candidate_id", "assembly_count", "complete_non_edge_instance_count"]].copy()
    numeric["assembly_count"] = pd.to_numeric(numeric["assembly_count"], errors="coerce")
    numeric["complete_non_edge_instance_count"] = pd.to_numeric(numeric["complete_non_edge_instance_count"], errors="coerce")
    s3_summary = pd.DataFrame([
        {"metric": "Genome-mapped candidates", "value": len(s3), "definition": "All rows in the genomic-support classification table."},
        {"metric": "Candidates with >=1 complete non-edge instance", "value": int((numeric["complete_non_edge_instance_count"] >= 1).sum()), "definition": "At least one complete non-edge genomic occurrence."},
        {"metric": "Principal genome-supported pool", "value": int(((numeric["complete_non_edge_instance_count"] >= 1) & (numeric["assembly_count"] >= 2)).sum()), "definition": "Complete non-edge support in >=2 assemblies."},
        {"metric": "Stringent recurrent subset", "value": int(((numeric["complete_non_edge_instance_count"] >= 1) & (numeric["assembly_count"] >= 10)).sum()), "definition": "Complete non-edge support in >=10 assemblies."},
    ])
    s3.to_csv(OUT / "Supplementary_Table_S3_candidate_genome_context_audit.tsv", sep="\t", index=False)
    s3_summary.to_csv(OUT / "Supplementary_Table_S3_audit_summary.tsv", sep="\t", index=False)
    write_book("Supplementary_Table_S3_candidate_genome_context_audit", {
        "S3_candidate_audit": s3,
        "S3_audit_summary": s3_summary,
    })

    # Table S4: join the 20-candidate evidence matrix to the exact experimental panel/MIC table.
    evidence = read_tsv("section_3_4/UNIFIED_20_EVIDENCE_TABLE.tsv")
    experimental = read_tsv("manuscript_tables/Table_1_Final_Experimental_Candidates_and_MIC.tsv")
    experimental = experimental.rename(columns={"Candidate ID": "candidate_id"})
    manifest = read_tsv("section_3_5/00_registry/candidate_manifest.tsv")
    s4 = evidence.merge(experimental, on="candidate_id", how="left", validate="one_to_one")
    s4 = s4.merge(manifest[["candidate_id", "amp_id"]], on="candidate_id", how="left", validate="one_to_one")

    coa_dir = ROOT / "section_3_5/00_registry/peptide_QC_and_reports/instrument_reports/biomatik_coa_docx"
    s4["synthesis_QC_document"] = s4["amp_id"].map(
        lambda amp: f"{amp}_Biomatik_COA.docx" if amp and (coa_dir / f"{amp}_Biomatik_COA.docx").exists() else "not found"
    )
    s4["synthesis_QC_status"] = s4["synthesis_QC_document"].map(
        lambda x: "Vendor COA present in project archive" if x != "not found" else "No COA file located"
    )

    mbc = pd.read_csv(
        ROOT / "section_3_5/04_analysis/MBC_Selected3Peptides_Analysis/Table_MBC_MIC_summary.csv",
        dtype=str,
        keep_default_na=False,
    ).rename(columns={"Peptide": "amp_id"})
    mbc["candidate_id"] = mbc["amp_id"].str.replace("AMP_", "CAND_", regex=False)
    mbc = mbc[["candidate_id", "amp_id", "Organism", "MIC (μM)", "Calculated MBC (μM)", "MBC/MIC", "Mean log10 reduction at 128 μM", "MBC basis", "Replicate/image status"]]
    mbc.to_csv(OUT / "Supplementary_Table_S4_preliminary_MBC_observations.tsv", sep="\t", index=False)

    qc_index = s4[["candidate_id", "amp_id", "synthesis_QC_document", "synthesis_QC_status"]].copy()
    s4.to_csv(OUT / "Supplementary_Table_S4_candidate_evidence_and_MIC.tsv", sep="\t", index=False)
    qc_index.to_csv(OUT / "Supplementary_Table_S4_synthesis_QC_index.tsv", sep="\t", index=False)
    write_book("Supplementary_Table_S4_candidate_evidence_and_experimental_data", {
        "S4_20_candidate_matrix": s4,
        "S4_preliminary_MBC": mbc,
        "S4_synthesis_QC_index": qc_index,
    })

    # Table S5: exact well-level MIC source data. Plate_Map_Trace preserves plate/well IDs.
    mic_raw_file = ROOT / "section_3_5/02_raw_data/MIC/MIC_8plates_BioTek_Epoch2_matrix_raw_export.xlsx"
    s5 = pd.read_excel(mic_raw_file, sheet_name="Plate_Map_Trace", dtype=str, keep_default_na=False)
    s5["well_class"] = s5["sample"].map(lambda x: "candidate" if str(x).startswith("CAND_") else "control")
    s5.to_csv(OUT / "Supplementary_Table_S5_MIC_well_level_OD600.tsv", sep="\t", index=False)
    write_book("Supplementary_Table_S5_MIC_well_level_OD600", {"S5_well_level_OD600": s5})

    # Table S6: MIC run record, calculation rule, control QC, summary, and selection logic.
    mic_doc = ROOT / "section_3_5/02_raw_data/MIC/MIC_experiment_documentation_package.xlsx"
    s6_run = pd.read_excel(mic_doc, sheet_name="Experiment_Run_Record", dtype=str, keep_default_na=False)
    s6_summary = pd.read_excel(mic_doc, sheet_name="Candidate_MIC_Summary", dtype=str, keep_default_na=False)
    s6_controls = pd.read_excel(mic_doc, sheet_name="Plate_Control_QC", dtype=str, keep_default_na=False)
    s6_rule = pd.read_excel(mic_doc, sheet_name="MIC_Calculation_Rule", dtype=str, keep_default_na=False)
    s6_selection = pd.read_excel(mic_doc, sheet_name="MBC_MIC_Selection", dtype=str, keep_default_na=False)
    s6_summary.to_csv(OUT / "Supplementary_Table_S6_MIC_summary.tsv", sep="\t", index=False)
    s6_controls.to_csv(OUT / "Supplementary_Table_S6_MIC_plate_controls.tsv", sep="\t", index=False)
    s6_rule.to_csv(OUT / "Supplementary_Table_S6_MIC_calculation_rule.tsv", sep="\t", index=False)
    write_book("Supplementary_Table_S6_MIC_QC_and_calling", {
        "S6_run_record": s6_run,
        "S6_MIC_summary": s6_summary,
        "S6_plate_control_QC": s6_controls,
        "S6_MIC_calling_rule": s6_rule,
        "S6_MBC_selection": s6_selection,
    })

    # Table S7: three-replicate MBC data.
    mbc_raw_file = ROOT / "section_3_5/02_raw_data/MBC/MBC_Optimized_3Rep.xlsx"
    s7_raw = pd.read_excel(mbc_raw_file, sheet_name="Raw_Counts_3Rep", dtype=str, keep_default_na=False)
    s7_calculated = pd.read_excel(mbc_raw_file, sheet_name="Calculated_CFU_MBC", dtype=str, keep_default_na=False)
    s7_summary = pd.read_excel(mbc_raw_file, sheet_name="MBC_Summary", dtype=str, keep_default_na=False)
    s7_images = pd.read_excel(mbc_raw_file, sheet_name="Image_Index", dtype=str, keep_default_na=False)
    s7_raw.to_csv(OUT / "Supplementary_Table_S7_MBC_raw_counts.tsv", sep="\t", index=False)
    s7_calculated.to_csv(OUT / "Supplementary_Table_S7_MBC_calculated_CFU.tsv", sep="\t", index=False)
    s7_images.to_csv(OUT / "Supplementary_Table_S7_MBC_image_index.tsv", sep="\t", index=False)
    write_book("Supplementary_Table_S7_MBC", {
        "S7_raw_counts_3Rep": s7_raw,
        "S7_calculated_CFU": s7_calculated,
        "S7_summary": s7_summary,
        "S7_image_index": s7_images,
    })

    # Table S8: peptide synthesis QC summary and the underlying HPLC/MS traces.
    qc_base = ROOT / "section_3_5/00_registry/peptide_QC_and_reports/instrument_reports/per_peptide_reports_final"
    s8 = pd.read_csv(qc_base / "20条候选肽_完整序列与HPLC_MS.tsv", sep="\t", dtype=str, keep_default_na=False)
    s8["candidate_id"] = "CAND_" + s8["candidate_id"].str.zfill(5)
    s8["amp_id"] = s8["candidate_id"].str.replace("CAND_", "AMP_", regex=False)
    s8["COA_document"] = s8["amp_id"].map(lambda x: f"{x}_Biomatik_COA.docx")
    s8["COA_status"] = "Vendor COA present in project archive"
    s8_trace = pd.read_csv(qc_base / "20_Candidates_HPLC_MS_Plot_Data_EN.tsv", sep="\t", dtype=str, keep_default_na=False)
    s8_trace["candidate_id"] = "CAND_" + s8_trace["candidate_id"].str.zfill(5)
    s8.to_csv(OUT / "Supplementary_Table_S8_peptide_synthesis_QC.tsv", sep="\t", index=False)
    s8_trace.to_csv(OUT / "Supplementary_Table_S8_HPLC_MS_trace_data.tsv", sep="\t", index=False)
    write_book("Supplementary_Table_S8_peptide_synthesis_QC", {
        "S8_QC_summary": s8,
        "S8_HPLC_MS_trace": s8_trace,
    })

    readme = """# Submission supplementary tables (S1–S8)\n\nAll values in this directory are copied or deterministically joined from audited project tables by `scripts/build_submission_supplementary_tables.py`. No values are imputed.\n\n| Table | Contents | Rows / sheets | Primary audited source |\n|---|---|---:|---|\n| Table S1 | Per-species genome and sORF resource statistics, plus aggregate statistics | 327 species + 1 summary row | `section_3_1/figures/Figure1_sampling_aware/data/` |\n| Table S2 | Model-support combinations and screening flow | 8 combinations + 6 flow rows | `section_3_2/data/` |\n| Table S3 | Candidate-level genomic occurrence/context audit and retention summary | 11,918 candidates + 4 summary rows | `section_3_3/01_master/` |\n| Table S4 | Complete 20-candidate evidence matrix, MIC annotations, MBC observations, and synthesis-QC index | 20 candidates + 6 MBC rows + 20 QC rows | `section_3_4/`, `manuscript_tables/`, `section_3_5/` |\n| Table S5 | Well-level MIC OD600 source data | 720 candidate wells | `section_3_5/02_raw_data/MIC/` |\n| Table S6 | MIC run record, plate controls, calculation rule, summary, and MBC-selection rationale | 5 sheets | `section_3_5/02_raw_data/MIC/` |\n| Table S7 | MBC three-replicate colony counts, calculated CFU, summary, and image index | 4 sheets | `section_3_5/02_raw_data/MBC/` |\n| Table S8 | Synthesis QC summary and HPLC/MS trace data | 20 peptides + 4,440 trace rows | `section_3_5/00_registry/peptide_QC_and_reports/` |\n\n**Important:** Table S7 reports MBC across all three biological replicates, all image-matched and verified.\n"""
    readme = readme.replace("720 candidate wells", "640 candidate wells + 80 control wells")
    (OUT / "README.md").write_text(readme, encoding="utf-8")
    print(f"[OK] Wrote submission tables to {OUT}")


if __name__ == "__main__":
    main()
```
