## Supplementary material for "Genome-context-aware discovery of antibacterial peptides from bacterial small open reading frames": Pathogen_sORF_AMP_Supplementary_Code: confirm_mbc_s7_metadata.py

```
#!/usr/bin/env python3
"""Align Table S7 wording with the confirmed three-biological-replicate record."""

from pathlib import Path
from openpyxl import load_workbook

ROOT = Path(__file__).resolve().parents[1]
TABLE = ROOT / "figures_generated" / "supplementary" / "tables" / "Supplementary_Table_S7_MBC.xlsx"


def main():
    wb = load_workbook(TABLE)
    ws = wb["S7_provisional_summary"]
    ws.title = "S7_MBC_summary"
    ws.cell(1, 5).value = "Replicate status"
    ws.cell(1, 6).value = "Interpretation"
    for row in range(2, ws.max_row + 1):
        ws.cell(row, 5).value = "n=3 independent biological replicates; Rep1/Rep2/Rep3 image-matched"
        ws.cell(row, 6).value = "Confirmed MBC endpoint using the prespecified ≥3-log10 reduction criterion"
    wb.save(TABLE)
    print(f"[OK] updated {TABLE}")


if __name__ == "__main__":
    main()
```
