## Supplementary material for "Genome-context-aware discovery of antibacterial peptides from bacterial small open reading frames": Pathogen_sORF_AMP_Supplementary_Code: plot_supplementary_figures.py

```
#!/usr/bin/env python3
"""
Supplementary figures S1-S6 (Nature style), based on
99_manuscript_preparation/04_key_figures_and_tables_for_manuscript.md
"""
import os
import numpy as np
import pandas as pd
import matplotlib
matplotlib.use("Agg")
import matplotlib.pyplot as plt
import matplotlib.image as mpimg

plt.rcParams.update({
    "font.family": "sans-serif",
    "font.sans-serif": ["Helvetica", "Arial", "DejaVu Sans"],
    "font.size": 7, "axes.labelsize": 7, "axes.titlesize": 7,
    "xtick.labelsize": 6.5, "ytick.labelsize": 6.5, "legend.fontsize": 6.5,
    "axes.linewidth": 0.6, "xtick.major.width": 0.6, "ytick.major.width": 0.6,
    "axes.spines.top": False, "axes.spines.right": False,
    "figure.dpi": 300, "savefig.dpi": 300, "savefig.bbox": "tight", "svg.fonttype": "none",
})
N_BLUE, N_RED, N_ORANGE, N_GREY, N_LIGHT_BLUE, N_GREEN = "#2c7bb6", "#d7191c", "#fdae61", "#969696", "#abd9e9", "#1a9641"

SC = "/media/qxl/EAGET/Pathogen_data/Pathogen_AMP_Mining/01_sorf_amp/02_screening_consensus"
RA = "/media/qxl/EAGET/Pathogen_data/Pathogen_AMP_Mining/01_sorf_amp/00_PROJECT_MASTER/results_section_audit"
OUT = os.path.join(RA, "figures_generated", "supplementary")
os.makedirs(OUT, exist_ok=True)

D03 = os.path.join(SC, "03_exact_sequence_distribution/figures")
D04 = os.path.join(SC, "04_sequence_cluster_robustness/figures")
D07 = os.path.join(SC, "07_pangenome_analysis/figures")
D07R = os.path.join(SC, "07_pangenome_analysis_refined/figures")
D08 = os.path.join(SC, "08_flagship_species/figures")
S5_MATRIX = os.path.join(SC, "05_exact_presence_absence/full_revalidation/near_ubiquitous_full_evidence_matrix.tsv")
S6_AUDIT = os.path.join(SC, "validation/outputs/cross_genus_identical_nt_audit.tsv")


def save(fig, name):
    fig.savefig(os.path.join(OUT, name + ".png"))
    fig.savefig(os.path.join(OUT, name + ".svg"))
    plt.close(fig)
    print(f"[OK] {name}")


def embed(ax, path, label, title=None):
    if os.path.exists(path):
        img = mpimg.imread(path)
        mask = np.any(img[..., :3] < .985, axis=2)
        yy, xx = np.where(mask)
        if len(xx):
            pad = max(8, int(min(img.shape[:2]) * .02))
            img = img[max(0, yy.min()-pad):min(img.shape[0], yy.max()+pad), max(0, xx.min()-pad):min(img.shape[1], xx.max()+pad)]
        ax.imshow(img)
    else:
        ax.text(0.5, 0.5, "missing", ha="center", va="center", fontsize=6, color="red")
    ax.axis("off")
    ax.text(-0.02, 1.02, label, transform=ax.transAxes, fontsize=8, fontweight="bold", va="bottom", ha="right")
    if title:
        ax.set_title(title, fontsize=6.5, loc="left", pad=2)


def S1():
    fig = plt.figure(figsize=(7.0, 6.5))
    gs = fig.add_gridspec(2, 2)
    embed(fig.add_subplot(gs[0, 0]), f"{D03}/fig2_species_count_distribution.png", "a")
    embed(fig.add_subplot(gs[0, 1]), f"{D03}/fig3_top15_species_absolute.png", "b")
    embed(fig.add_subplot(gs[1, 0]), f"{D03}/fig4_top15_species_normalized.png", "c")
    embed(fig.add_subplot(gs[1, 1]), f"{D03}/fig7_frequency_categories.png", "d")
    fig.suptitle("Supplementary Figure S1 | Assembly- and species-level exact-sequence distribution", y=1.0, fontsize=9, fontweight="bold")
    save(fig, "Supp_FigS1_exact_distribution")


def S2():
    fig = plt.figure(figsize=(7.0, 3.0))
    gs = fig.add_gridspec(1, 2)
    embed(fig.add_subplot(gs[0, 0]), f"{D04}/fig1_coverage_sensitivity.png", "a")
    embed(fig.add_subplot(gs[0, 1]), f"{D04}/fig2_length_analysis.png", "b")
    fig.suptitle("Supplementary Figure S2 | Clustering robustness and positive-control validation", y=1.02, fontsize=9, fontweight="bold")
    save(fig, "Supp_FigS2_cluster_robustness")


def S3():
    fig = plt.figure(figsize=(7.0, 6.5))
    gs = fig.add_gridspec(2, 2)
    embed(fig.add_subplot(gs[0, 0]), f"{D07}/fig1_gram_comparison.png", "a")
    embed(fig.add_subplot(gs[0, 1]), f"{D07}/fig2_eskape_comparison.png", "b")
    embed(fig.add_subplot(gs[1, 0]), f"{D07R}/fig1_cliffs_delta_gram.png", "c")
    embed(fig.add_subplot(gs[1, 1]), f"{D07R}/fig2_zero_inflation.png", "d")
    fig.suptitle("Supplementary Figure S3 | Pangenome comparison and effect-size refinement", y=1.0, fontsize=9, fontweight="bold")
    save(fig, "Supp_FigS3_pangenome_refinement")


def S4():
    fig = plt.figure(figsize=(7.0, 9.0))
    gs = fig.add_gridspec(3, 2)
    # Only include species with an actual heatmap export; never show a "missing" placeholder.
    sp = ["Klebsiella_pneumoniae", "Staphylococcus_aureus", "Mycobacterium_tuberculosis",
          "Bacillus_pumilus", "Burkholderia_pseudomallei", "Stenotrophomonas_maltophilia"]
    display = {
        "Klebsiella_pneumoniae": "Klebsiella pneumoniae",
        "Staphylococcus_aureus": "Staphylococcus aureus",
        "Mycobacterium_tuberculosis": "Mycobacterium tuberculosis",
        "Bacillus_pumilus": "Bacillus pumilus",
        "Burkholderia_pseudomallei": "Burkholderia pseudomallei",
        "Stenotrophomonas_maltophilia": "Stenotrophomonas maltophilia",
    }
    for i, s in enumerate(sp):
        r, c = divmod(i, 2)
        embed(fig.add_subplot(gs[r, c]), f"{D08}/heatmap_{s}.png", chr(97 + i), display[s])
    fig.suptitle("Supplementary Figure S4 | Strain-level candidate heatmaps across representative species", y=1.0, fontsize=9, fontweight="bold")
    save(fig, "Supplementary_FigS4_flagship_species_heatmaps_optimized")


def S5():
    df = pd.read_csv(S5_MATRIX, sep="\t")
    fig, ax = plt.subplots(figsize=(7.0, 3.0))
    ax.axis("off")
    cols = ["candidate_id", "evidence_completeness", "experimental_priority"]
    data = df[cols].values.tolist()
    header = ["Candidate", "Evidence completeness", "Priority"]
    tbl = ax.table(cellText=data, colLabels=header, loc="center", cellLoc="left")
    tbl.auto_set_font_size(False)
    tbl.set_fontsize(6.5)
    tbl.scale(1, 1.4)
    for (r, c), cell in tbl.get_celld().items():
        cell.set_edgecolor(N_GREY)
        cell.set_linewidth(0.4)
        if r == 0:
            cell.set_facecolor(N_LIGHT_BLUE)
            cell.set_text_props(fontweight="bold")
    ax.set_title(f"Supplementary Figure S5 | Revalidation of near-ubiquitous candidates (n={len(df)})", loc="left", fontsize=9, fontweight="bold")
    save(fig, "Supp_FigS5_near_ubiquitous_revalidation")


def S6():
    df = pd.read_csv(S6_AUDIT, sep="\t")
    fig = plt.figure(figsize=(7.0, 3.0))
    gs = fig.add_gridspec(1, 3)
    # a: classification
    ax = fig.add_subplot(gs[0, 0])
    vc = df["classification"].value_counts()
    ax.bar(vc.index, vc.values, color=[N_BLUE, N_GREY][:len(vc)], width=0.6, linewidth=0)
    for i, v in enumerate(vc.values):
        ax.text(i, v, str(v), ha="center", va="bottom", fontsize=6)
    ax.set_xlabel("Classification")
    ax.set_ylabel("Cross-genus candidates")
    ax.text(-0.02, 1.02, "a", transform=ax.transAxes, fontsize=8, fontweight="bold")
    # b: n_species distribution
    ax = fig.add_subplot(gs[0, 1])
    ax.hist(df["n_species"], bins=np.arange(2, 8) - 0.5, color=N_ORANGE, edgecolor="white", linewidth=0.3)
    ax.set_xlabel("Number of species")
    ax.set_ylabel("Candidates")
    ax.text(-0.02, 1.02, "b", transform=ax.transAxes, fontsize=8, fontweight="bold")
    # c: nt_identical vs plasmid_signal
    ax = fig.add_subplot(gs[0, 2])
    nt = df["nt_identical"].astype(bool).sum()
    pl = df["plasmid_signal"].astype(bool).sum()
    ax.bar(["nt-identical", "plasmid signal"], [nt, pl], color=[N_GREEN, N_RED], width=0.5, linewidth=0)
    for i, v in enumerate([nt, pl]):
        ax.text(i, v, str(v), ha="center", va="bottom", fontsize=6)
    ax.set_ylabel("Candidates")
    ax.text(-0.02, 1.02, "c", transform=ax.transAxes, fontsize=8, fontweight="bold")
    fig.suptitle("Supplementary Figure S6 | Cross-species extent and artifact audit", y=1.02, fontsize=9, fontweight="bold")
    save(fig, "Supp_FigS6_cross_species_audit")


def main():
    print("=== Supplementary figures S1-S6 ===")
    S1(); S2(); S3(); S4(); S5(); S6()
    print(f"=== Done: {OUT} ===")


if __name__ == "__main__":
    main()
```
