## Supplementary material for "Genome-context-aware discovery of antibacterial peptides from bacterial small open reading frames": Pathogen_sORF_AMP_Supplementary_Code: reconcile_new_mic_files_to_old_source.py

```
"""Rebuild the three MIC companion workbooks from the authoritative source.

The source is OD600_dataset.xlsx, version with 10 valid A-H / 1-12 plates.
No OD600 value, plate-map entry, or MIC summary value is altered.
"""

from pathlib import Path
from collections import defaultdict
from openpyxl import Workbook, load_workbook


BASE = Path(__file__).resolve().parents[1]
MIC = BASE / "02_raw_data" / "MIC"
SOURCE = MIC / "OD600_dataset.xlsx"
RAW_OUT = MIC / "MIC_8plates_BioTek_Epoch2_matrix_raw_export.xlsx"
MAP_OUT = MIC / "MIC_8plates_Plate_Map.xlsx"
DOC_OUT = MIC / "MIC_experiment_documentation_package.xlsx"
PLATES = [(prefix, strain, number) for prefix, strain in (("EC", "E.coli"), ("SA", "S.aureus")) for number in range(1, 6)]


def map_sheet_name(prefix, number):
    return f"{prefix}_Plate{number:03d}_Plate_Map"


def raw_sheet_name(prefix, number):
    return f"{prefix}_Plate{number:03d}_Raw_OD600"


def raw_matrix(source, prefix, number):
    ws = source[raw_sheet_name(prefix, number)]
    return {
        f"{ws.cell(row=r, column=1).value}{ws.cell(row=5, column=c).value}": ws.cell(row=r, column=c).value
        for r in range(6, 14)
        for c in range(2, 14)
    }


def map_rows(source, prefix, number):
    ws = source[map_sheet_name(prefix, number)]
    return [tuple(row) for row in ws.iter_rows(min_row=2, values_only=True) if row[0] is not None]


def write_source_metadata(ws, info):
    ws.append(["Parameter", "Value"])
    ws.append(["Authoritative_source", SOURCE.name])
    ws.append(["Transformation", "Copied directly from the authoritative source; no OD600, map, or MIC value was changed."])
    for key, value in info.items():
        ws.append([key, value])


def build_raw_export(source, info, summary):
    out = Workbook()
    out.remove(out.active)
    trace = out.create_sheet("Plate_Map_Trace")
    trace.append(["plate_id", "well", "sample", "strain", "concentration", "technical_replicate", "od600"])
    for prefix, strain, number in PLATES:
        name = f"{prefix}_Plate{number:03d}_RawMatrix"
        ws = out.create_sheet(name)
        ws.append([f"Exact OD600 matrix copied from {SOURCE.name}: {raw_sheet_name(prefix, number)}"])
        ws.append(["Read", "Absorbance OD600", "Plate", f"{prefix}_Plate{number:03d}"])
        ws.append([])
        ws.append(["Well", *range(1, 13)])
        matrix = raw_matrix(source, prefix, number)
        for row in "ABCDEFGH":
            ws.append([row, *[matrix[f"{row}{column}"] for column in range(1, 13)]])
        for well, sample, mapped_strain, concentration, replicate in map_rows(source, prefix, number):
            trace.append([f"{prefix}_Plate{number:03d}", well, sample, mapped_strain, concentration, replicate, matrix[well]])
    ws = out.create_sheet("MIC_Summary")
    ws.append(["Candidate_ID", "E_coli_MIC_uM", "S_aureus_MIC_uM"])
    for row in summary:
        ws.append(list(row))
    write_source_metadata(out.create_sheet("Source_Metadata"), info)
    out.save(RAW_OUT)


def build_plate_map(source):
    out = Workbook()
    out.remove(out.active)
    overview = out.create_sheet("Plate_Summary")
    overview.append(["plate_id", "strain", "mapped_wells", "candidate_wells", "control_wells", "unmeasured_wells", "source"])
    for prefix, strain, number in PLATES:
        matrix = raw_matrix(source, prefix, number)
        entries = map_rows(source, prefix, number)
        ws = out.create_sheet(map_sheet_name(prefix, number))
        ws.append(["Well", "Sample", "Strain", "Concentration", "Replicate", "OD600"])
        for well, sample, mapped_strain, concentration, replicate in entries:
            ws.append([well, sample, mapped_strain, concentration, replicate, matrix[well]])
        candidate_wells = sum(1 for row in entries if isinstance(row[1], str) and row[1].startswith("CAND_"))
        overview.append([f"{prefix}_Plate{number:03d}", strain, len(entries), candidate_wells, len(entries) - candidate_wells, 96 - len(entries), SOURCE.name])
    out.save(MAP_OUT)


def control_values(source, prefix, number):
    matrix = raw_matrix(source, prefix, number)
    controls = defaultdict(list)
    for well, sample, *_ in map_rows(source, prefix, number):
        if not (isinstance(sample, str) and sample.startswith("CAND_")):
            controls[sample].append(matrix[well])
    return {name: values for name, values in controls.items()}


def build_documentation(source, info, summary):
    out = Workbook()
    run = out.active
    run.title = "Experiment_Run_Record"
    write_source_metadata(run, info)
    registry = out.create_sheet("Candidate_MIC_Summary")
    registry.append(["Candidate_ID", "E_coli_MIC_uM", "S_aureus_MIC_uM"])
    for row in summary:
        registry.append(list(row))
    qc = out.create_sheet("Plate_Control_QC")
    qc.append(["plate_id", "control_type", "replicate_1_od600", "replicate_2_od600", "mean_od600"])
    for prefix, _, number in PLATES:
        for control, values in control_values(source, prefix, number).items():
            qc.append([f"{prefix}_Plate{number:03d}", control, values[0], values[1], sum(values) / len(values)])
    method = out.create_sheet("MIC_Calculation_Rule")
    method.append(["Item", "Definition"])
    method.append(["Input", "Well-level OD600 + matching plate map"])
    method.append(["Blank correction", "OD_sample - mean(Blank controls)"])
    method.append(["Growth reference", "mean(Growth control) - mean(Blank controls)"])
    method.append(["Inhibition", "1 - (sample corrected / growth reference)"])
    method.append(["MIC threshold", ">=90% inhibition using the mean of two technical replicates"])
    selection = out.create_sheet("MBC_MIC_Selection")
    selection.append(["Role", "Candidate_ID", "E_coli_MIC_uM", "S_aureus_MIC_uM", "Rationale"])
    values = {row[0]: row[1:] for row in summary}
    for role, candidate, rationale in [
        ("Broad-spectrum", "CAND_04141", "Lowest worst-case MIC across both organisms"),
        ("Gram-negative priority", "CAND_07825", "Lowest remaining E. coli MIC; stronger S. aureus MIC than the E. coli tie"),
        ("Gram-positive priority", "CAND_04265", "Lowest remaining S. aureus MIC with stronger E. coli MIC than the S. aureus tie"),
    ]:
        ec, sa = values[candidate]
        selection.append([role, candidate, ec, sa, rationale])
    out.save(DOC_OUT)


def main():
    source = load_workbook(SOURCE, data_only=True, read_only=True)
    info = {row[0]: row[1] for row in source["Experiment_Info"].iter_rows(min_row=2, values_only=True)}
    summary = list(source["MIC_Summary"].iter_rows(min_row=2, values_only=True))
    build_raw_export(source, info, summary)
    build_plate_map(source)
    build_documentation(source, info, summary)


if __name__ == "__main__":
    main()
```
